# Microbiomes in drinking water treatment and distribution: a meta-analysis from source to tap

**DOI:** 10.1101/2021.08.30.457654

**Authors:** Claire Thom, Cindy J Smith, Graeme Moore, Paul Weir, Umer Z Ijaz

## Abstract

A meta-analysis of existing and available Illumina 16S rRNA datasets from drinking water source, treatment and drinking water distribution systems (DWDS) were collated to compare changes in abundance and diversity throughout. Samples from bulk water and biofilm were used to assess principles governing microbial community assembly and the value of amplicon sequencing to water utilities. Individual phyla relationships were explored to identify competitive or synergistic factors governing DWDS microbiomes. The relative importance of stochasticity in the assembly of the DWDS microbiome was considered to identify the significance of source and treatment in determining communities in DWDS. Treatment of water significantly reduces overall species abundance and richness, with chlorination of water providing the most impact to individual taxa relationships. The assembly of microbial communities in the bulk water of the source, primary treatment process and DWDS is governed by more stochastic processes, as is the DWDS biofilm. DWDS biofilm is significantly different from bulk water in terms of local contribution to beta diversity, type and abundance of taxa present. Water immediately post chlorination has a more deterministic microbial assembly, highlighting the significance of this process in changing the microbiome, although elevated levels of stochasticity in DWDS samples suggest that this may not be the case at customer taps. 16S rRNA sequencing is becoming more routine, and may have several uses for water utilities, including: detection and risk assessment of potential pathogens such as those within the genera of *Legionella* and *Mycobacterium;* assessing the risk of nitrification in DWDS; providing improved indicators of process performance and monitoring for significant changes in the microbial community to detect contamination. Combining this with quantitative methods like flow cytometry will allow a greater depth of understanding of the DWDS microbiome.

## 1. Introduction

The safety of drinking water supplies is of paramount importance for public health. Water utilities are responsible for the treatment and delivery of potable water. While treatment is highly effective at removing traditional faecal indicator organisms, the microbial challenge remains significant as water-borne disease outbreaks associated with drinking water distribution systems (DWDS) still have significant public health implications, which may not correlate with traditional water quality metrics (Fricker and Eldred, 2009; Jalava et al., 2014; Leclerc et al., 2001; Payment and Locas, 2011; Saxena et al., 2015). For example, in 1998, an outbreak of campylobacteriosis in northern Finland was not detected by routine water quality samples. Contamination occurred during a routine repair to a mains pipe, leading to acute gastroenteritis in over 200 people (Kuusi et al., 2005).

Water treatment has traditionally been a 3-stage process: Coagulation of colloidal material using a metallic salt; subsequent filtration through rapid gravity sand filters and disinfection using a chlorine-based biocide. This remains the most common method of treatment in many countries. There are several other treatment strategies that are now used, which satisfy drinking water regulations, including: slow sand filtration, biological filtration, ozonation and membrane filtration. Filtration and disinfection are the primary barriers to the presence of harmful pathogens in drinking water.

Water quality regulations on the microbial safety of DWDS focus on the likelihood of faecal contamination using the presence of coliform bacteria as a surrogate for the wide range of pathogens potentially present in faeces. These are measured using culture-based tests that isolate and enumerate coliforms and *E. coli* specifically. These methods have been broadly unchanged for over 100 years. Compliance with these metrics is high in the UK (>99.9%, (DWI, 2020), although isolated, sporadic, and low-level total coliform detections remain a problem for utilities, often without an attributable cause. These indicators are now known to be problematic for a few reasons: more than 99% of bacteria are unculturable (Hahn et al., 2019; Rappé and Giovannoni, 2003); there are also emerging non-faecal pathogens in drinking water, e.g., *Mycobacterium* and *Legionella* species; and the correlation between total coliforms and other pathogenic indicators is poor (Cabral, 2010; Payment and Locas, 2011; Savichtcheva and Okabe, 2006). Moreover, routine culture tests assess a small volume (~100 mL), and a confirmed result takes over two days, meaning that poor quality water will have already passed into the DWDS well before a result is returned. Quantitative polymerase chain reaction (qPCR) methods are capable of identification of specific pathogens or target organisms within a few hours but are limited in the amount of information they give about the overall microbiome. There is therefore a need for additional high-throughput methods of microbial characterisation to assess the diversity of microbial communities across space and time. These approaches will need to move beyond the limitations of day-to-day testing for specific pathogenic microbes, while assessing changes in the microbiome at a macro level over a long-term period to manage DWDS proactively.

16S rRNA amplicon sequencing technology can be used to characterise and identify microbial communities in DWDS across space and time. While the taxonomic resolution that can be achieved depends on the 16S rRNA hypervariable region sequenced and the type and abundance of the taxa detected (Fox and Reid-Bayliss, 2014), the approach has been widely applied in academia since circa 2010 to explore microbial communities of drinking water to both assess diversity and identify groups of microbes of concern to public health. This method has broad advantages over qPCR in the large amount of taxonomic information that it provides, although it is non-quantitative and unable to determine cell viability. Most 16S rRNA studies are discrete, across a single or few DWDS within a geographical area. They also tend to focus on one part of a system, e.g., source waters; efficacy of treatment processes; variations of biofilm communities in space, time, or operating conditions within a pipe distribution network; influences of domestic plumbing arrangements, or differences between the bulk water and biofilm communities (Ahmed et al., 2015; Bautista-de los Santos et al., 2016; Bautista-De los Santos et al., 2016; Chan et al., 2019; De Vera et al., 2018; Douterelo et al., 2019, 2017; Fish and Boxall, 2018; Gerrity et al., 2018; Ghaju Shrestha et al., 2017; Gülay et al., 2016; Han et al., 2020; Hou et al., 2018; Lautenschlager et al., 2014; Liu et al., 2017b; Lührig et al., 2015; McCoy and VanBriesen, 2012; Potgieter et al., 2018; Potgieter and Pinto, 2019; Prest et al., 2014; Shaw et al., 2015; Uyaguari-Diaz et al., 2019; Vierheilig et al., 2015; Vignola et al., 2018; Wan et al., 2019; Wang et al., 2014, 2013; Wolf-Baca and Piekarska, 2020; Wu et al., 2014; Zhu et al., 2019). While several of these studies have provided new insight into drinking water microbiomes, they tend to be descriptive and not predictive. Add to this variability in the methods used to construct amplicon libraries (e.g. DNA extraction, 16S rRNA hypervariable region and sequencing platform) and the complex system-specific nature of DWDS and building a general understanding of how drinking water microbial communities change from source to tap becomes difficult.

16S rRNA studies have shown the treatment of drinking water, in general, reduces the abundance and diversity of micro-organisms present, yet a diverse microbiome remains in potable water, including genera containing pathogenic species. The drinking water microbiome at customer taps may be influenced by a range of factors, including: source water, treatment, flow conditions and DWDS biofilms. Water treatment has been proposed to have a deterministic effect, selecting microbes that survive filtration and disinfection processes (Pinto *et al*., 2012; Lin *et al*., 2014). This effect is likely to reduce with distance and time from treatment, where biofilm growth and disturbance become more prominent. At this point, stochastic (random) effects may be more likely to govern the assembly of microbial communities due to the complexity of the local environments. The importance of stochastic factors like birth, death and immigration are known to be important in shaping many prokaryotic communities (Sloan et al., 2006). Thus, drinking water microbiomes are dynamic through treatment, time, and location. To aid water utilities to deliver safe potable water, a deeper understanding of these changes, consequences, and impact on both the microbiome and the prevalence of pathogens is needed.

### Aims and objectives

Here we present a meta-analysis of 16S rRNA studies from source to tap to explore global distribution and commonalities in the drinking water microbiomes. We further consider the contribution and potential of 16S rRNA amplicon sequencing as an analytical tool for water utilities. Research has suggested that 16S rRNA amplicon sequencing is beneficial in assessing risk to public health in DWDS, although there are many areas for further investigation to understand the implications of the results (Vierheilig et al., 2015). The specific aims of this meta-analysis were to identify commonalities in DWDS microbiomes across the world, which can be used to further understanding of water quality for utilities; to understand the relative importance of the deterministic effects of source and treatment on the microbiomes of DWDS; and explore key relationships between phyla present.

## 2. Methods

### 2.1 Data Gathering

A search for all papers since 2010 using the following terms: “16S rRNA” and “Water” was carried out using Scopus. This search returned 176 results. Each result was individually assessed to ascertain its relevance to this meta-analysis. Only studies using Illumina MiSeq or HiSeq^®^ platforms were included to minimise the different errors and biases associated with alternative sequencing platforms such as Nanopore^®^, Ion Torrent^®^ or older technologies used before 2010 such as Pyrosequencing (D’Amore et al., 2016; Schirmer et al., 2015). After this manual filter, 44 studies remained and were checked to ascertain sequence data availability. 26 studies had publicly available raw sequence data. For the remainder, data was requested. Only one study responded. A list of the papers used in the analysis can be found in the supplementary information (Supplementary Information 1). All raw data downloads used the SRA Toolkit provided by NCBI, except for one study from QIITA. Metadata for samples from NCBI’s Run Selector included: sequencing platform; the hypervariable region of the 16S rRNA gene sequenced; sample I.D.; sample date and time; and geolocation. Other relevant metadata was recorded from the published research: sample location, disinfection type (if applicable), and whether the sample was from bulk water or biofilm. Before processing, studies were grouped by the hypervariable region of the 16S rRNA gene sequenced. All studies included in this meta-analysis and relevant sample information are listed in Supplementary Information 1. In total 27 studies, with 1750 samples, from over 50 different DWDS were compared.

### 2.2 Sequence Processing

QIIME2 processed collated amplicon sequences for each platform and hypervariable V-region in Earth Microbiome Project Paired-end Sequencing Format (.fastq) and was used to generate Amplicon Sequencing Variants (ASVs). QIIME2 improves QIIME1 in terms of quality control of sequences using DADA2 and Deblur software, both of which were employed here. To provide enough overlap of forward and reverse reads to facilitate paired-end reads, DADA2 was employed where amplicons were <250bp long and the quality score was >20. For amplicons spanning multiple V regions, DEBLUR commands allowed for the pairing of longer amplicons without significant loss of sequence length, as an explicit threshold is not required. Output alpha diversity profiles may be significantly different when using different denoising software to generate ASVs (Nearing et al., 2018), so runs of DEBLUR and DADA2 were carried out for all regions and platforms and compared. The final analyses generated 3.32 × 10^8^ demultiplexed reads in total from 2098 samples.

To identify the best taxonomic assignment, abundance tables and phylogenetic tree files from QIIME2 had taxonomy assigned using three approaches. These were: Naïve Bayesian Classification system (NBC), Bayesian Least Common Ancestor (BLCA) approach (using SILVA138 database), and the TaxAss database. TaxAss uses SILVA to generate a first pass of taxonomic assignment then a curated database of freshwater sequences to assign the remainder of ASVs. BLCA and SILVA138 were selected for downstream statistical processing as this method provided the highest level of taxonomic recovery to the genus level (Supplementary Information 2). Finally, ASVs from all V regions were collated together in a single abundance table. ASVs without sequence-level resolution were removed so that full-length 16S rRNA sequences could be obtained for all sequences as per the method used by Keating et al (Keating et al., 2020). Of the 1.27 x10^8^ sequences originally classified by SILVA138 from the individual datasets, 1.08 x10^8^ were classified to sequence level, that is 84.89%. Of these sequences, 22574 were unique taxa. Generation of the final phylogenetic tree and abundance table with taxonomy was again processed in QIIME2.

The collation methodology applied in this meta-analysis is limited in that only full-length sequences already present in the SILVA database are included. This may lead to bias towards these sequences. In total from our data base, 15.11% of sequences were not present in SILVA with sequence-level resolution. However, without the collation strategy, 16S rRNA amplicon sequencing datasets of different hypervariable regions cannot be pooled and compared. Despite the loss of these sequences, the overall beta diversity patterns present in the uncollated and collated datasets were largely preserved as shown by Mantel and Procrustes analyses (Supplementary information 3). Only one region/platform correlated poorly (MiSeq and V3), due to differences in the abundances of 2 uncultured sequences between the datasets, with the rest of the taxa being identical. This means that the between sample diversity patterns for collated and uncollated samples were highly similar, indicating that the removed sequences were in low abundance and unlikely to disproportionally affect the results.

### 2.3 Statistical Analyses

The collated abundance table with taxonomy, phylogenetic tree, and metadata was then processed using the microbiome seq packages (R). Meta-sample groupings defined the sample location in the treatment and distribution process and if the sample originated from biofilm or bulk water. Samples from sediments and wastewater streams were removed at this stage, giving a final sample number of 1750. Shannon and Richness indexes were calculated for each meta-grouping to estimate alpha diversity (diversity within a sample). Core microbiome analysis of the collated datasets was carried out in the Bioconductor package, using an absolute detection method and a minimum prevalence of 85% for all groups except DWDS and Untreated water groups. These groups had significantly more samples and required a higher threshold of 95% (Lahti et al., 2017).

Beta diversity (or between-sample diversity) metrics were more complicated to assess, given the substantial number of samples in the final analysis (n=1750), their varying environments as well as spatial and temporal locations. Instead, calculation of Local Contribution to Beta Diversity (LCBD) for each group was made (Legendre and De Cáceres, 2013). The Nearest Taxon Index (NTI) and Net Relatedness Index (NRI) from the Picante package in R (http://kembellab.ca/r-workshop/biodivR/SK_Biodiversity_R.html) were used to quantify Environmental filtering on community assembly.

To estimate the relative impacts of ecological determinism and stochasticity on the assembly of the curated microbiomes a null modelling approach was adopted using a general framework defined by Ning et al. 2019, using simulated bacterial communities and further applied to real communities (Nikolova et al., 2021; Ning et al., 2019; Trego et al., 2021). Normalised and Modified Stochasticity Ratios (NST and MST) were calculated for all groups within the dataset. In this approach, deterministic processes are expected to drive species more similar or dissimilar than the null expectation. NST values are quantified from the difference between the expected similarity and dissimilarity and the actual values. Higher NST/MST (above 0.5) values indicate more deterministic factors influencing community assembly, lower values (below 0.5) indicate more stochastic influences.

Patterns in beta diversity may not be continual, as multiple relationships may affect an organism at a specific time or place. Therefore, a new methodology by Golovko *et al*. (2020) was employed using boolean patterns to assess relationships between individual ASVs in all meta-sample groups. This uses a pattern-specific method to identify relationships between 2 ASVs at a defined threshold, including one-way relationships, co-occurrence, and co-exclusion. This method can also quantify 3-dimensional relationships between ASVs. These are categorised as: all alone (type 1 co-exclusion); exclusion of ASV1 by ASV2 and 3 (type 2 co-exclusion); presence of ASV2 and ASV3 if ASV1 is present, and finally, all three together. This method was applied to the ASVs in the dataset to identify any significant relationships at a phyla level.

## 3. Results

### 3.1 Taxonomic Profile

Sequence-level classification was resolved for a total of 22754 ASVs. The 25 most abundant genera are shown in Figure 2. 1. In the final analysis, 111 samples, were from DWDS bulk water, the largest meta-sample group. Bulk water from different DWDS, as expected, was variable with differences in the abundances of the top 25 genera. However, there does appear to be some commonalities in taxa among DWDS with the same disinfectant residual: *Nitrosomonas and Pseudomonas* were more abundant in systems using a chloraminated residual. Pathogenic microbes such as *Mycobacterium* were common in both chlorinated and chloraminated systems. Biofilm samples in distribution were less numerous (n=159) and had a much higher taxonomic diversity than the bulk water. *Pseudomonas* was common in many samples in both chlorinated and chloraminated biofilms, but less so in those with no disinfectant residual.

**Figure 1:**
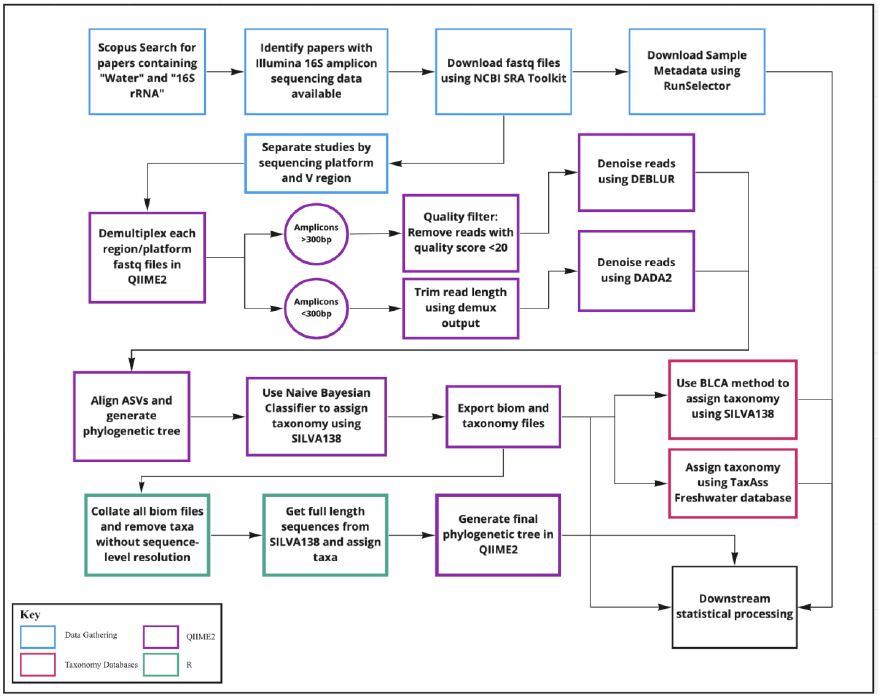
Methods: An overview of the methodologies used to generate Amplicon Sequencing Variants (ASVs) for this meta-analysis.

**Figure 2:**
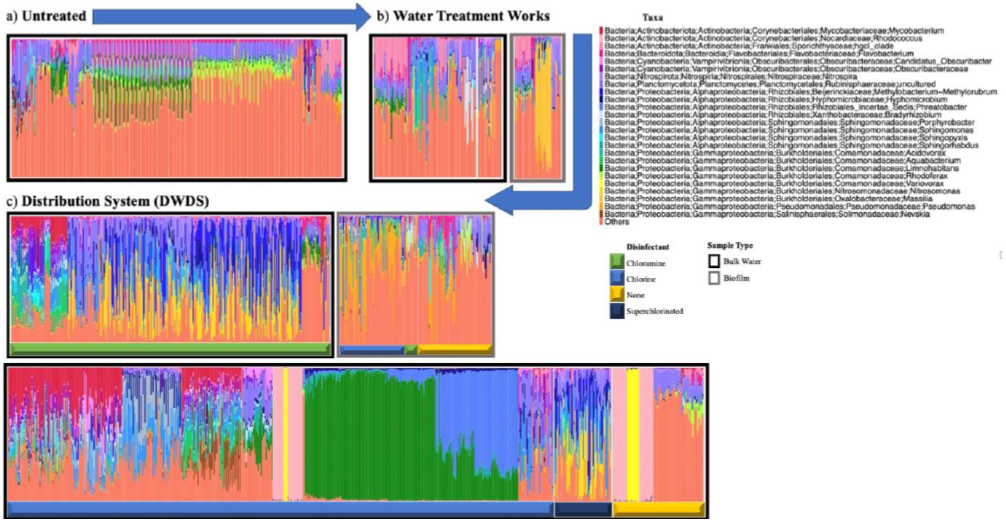
Taxa Profiles: Proportion of most abundant genera for all samples in this meta-analysis, grouped by: A untreated water; B water and biofilm isolated throughout treatment processes; C biofilm and bulk water sampled from distribution pipes. Where appropriate, samples have been colour-coded by disinfectant residual: Free chlorine (blue); Chloraminated (green); None (yellow); Superchlorinated (dark blue). Samples are further grouped by study.

Samples from water sources and treatment systems made up a much smaller proportion of the dataset and differed from DWDS in terms of the most abundant taxa. Again, the most common genera in DWDS were less abundant in source and treatment, except *Nitrospira*, which was more abundant throughout treatment than in distribution. A member of *Burkholderiales, Limnohabitans* was also present throughout treatment and highly abundant in one bulk water study in the distribution. Source water samples were generally from surface waters, although a small proportion came from groundwater. Several untreated water samples show similar taxonomic profiles to each other. Globally all untreated water samples were highly diverse in comparison to treated water.

### 3.2 Alpha Diversity & Core Microbiome Analysis of Collated Datasets

The amount of diversity within each sample, or alpha diversity, can be seen in Figure 3(C). Across the different sample groups, the within-sample richness values were significantly different. The highest degree of sequence diversity in terms of richness and Shannon index values came from untreated water. A reduction in these values was evident in the treatment and DWDS groups, in biofilm and bulk water. Biofilm samples have elevated Shannon values compared to bulk water, although richness was similar. Core microbiome analysis of this collated dataset proposed several prevalent taxa within more than one sample group, although the overall taxa prevalence was reduced in distribution samples. *Pseudomonas* was the most commonly abundant taxa and was present at all stages of water treatment and in DWDS biofilm. *Nitrospira* was prevalent within water treatment works, bulk water and biofilm. *Legionella* was abundant in bulk water only, in untreated and in treatment samples. The only taxa prevalent throughout all bulk water and biofilm groups was an uncultured *Rubinisphaeraceae*.

**Figure 3:**
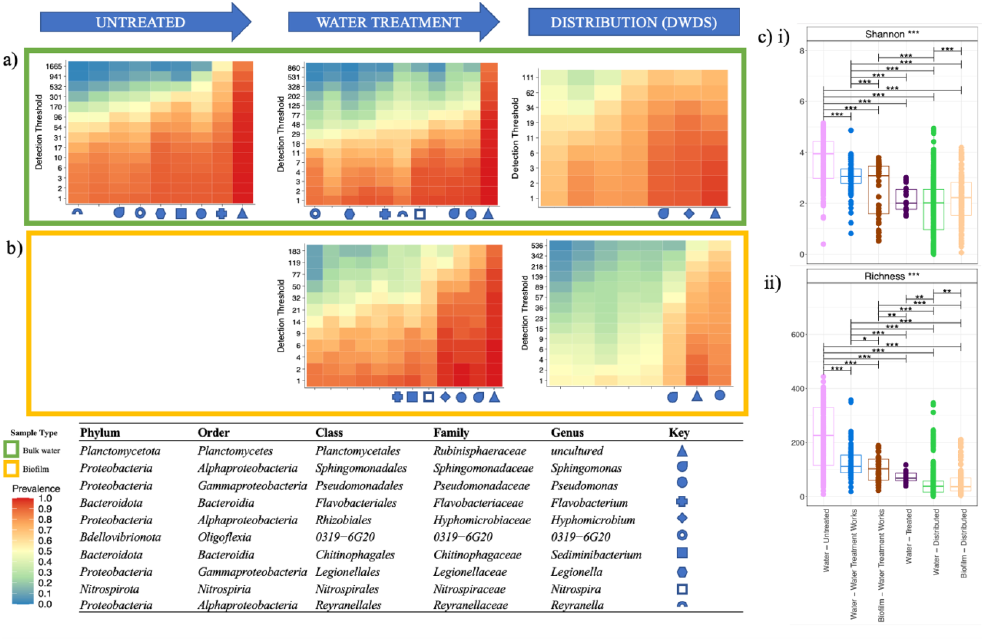
Core Microbiome Analysis: Heatmaps of the different meta-sample groups from source through treatment and distribution for bulk water (a)) and for biofilm (b)). Minimum prevalence was set at 0.85 for all groups except Distributed Bulk Water and Untreated water, set at 0.95 due to the high number of samples in those groups. c) the alpha diversity of the various meta-sample groups, displaying i) Shannon values and ii) Richness.

### 3.3 Local Contribution to Beta Diversity and Environmental Filtering

Due to the unequal data classes and high degrees of spatial and temporal variation, estimates of beta diversity used Local Contribution to Beta diversity (LCBD) for all meta-sample groups (Figure 4) rather than a direct measure. LCBD values were only above the significance threshold for two categories when calculated using Unifrac distance: untreated water and distribution biofilm. For Bray-Curtis, all groups had greater than the calculated threshold (0.00057) LCBD except distribution water, with DWDS biofilm and treated water samples having the highest values. The untreated and water treatment works groups were very close to the significance threshold. Unifrac shows a pattern of reducing LCBD in bulk water throughout treatment and distribution, whereas biofilm groups increase again in DWDS samples. Bray-Curtis measures show a pattern of increasing LCBD in bulk water samples from untreated to treated water, which then reduces in DWDS. LCBD in biofilm samples increased throughout treatment to a maximum in DWDS. NTI and NRI values for the meta-sample groups were similar except for untreated water, which was the only category with values <0, indicating the taxa present are more dissimilar than in the other categories. Biofilm in DWDS had a higher NTI than that of bulk water indicating that species relatedness is higher in biofilm.

**Figure 4:**
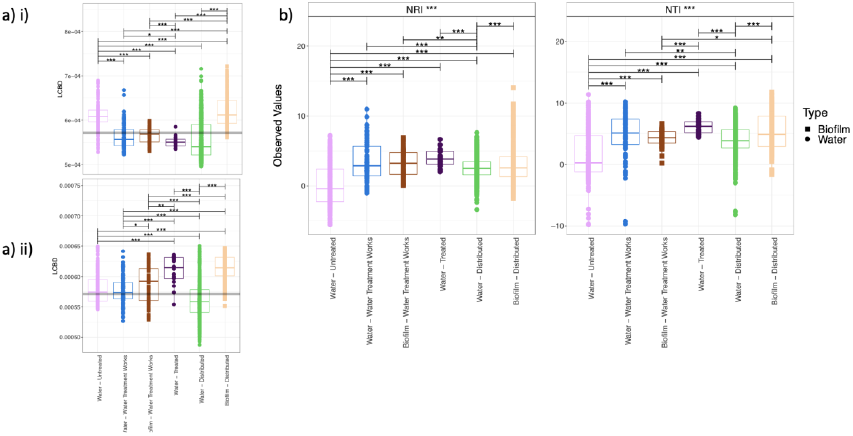
Local contribution to beta diversity (LCBD): a) LCBD values for all meta-sample groups using Unifrac (i)) and Bray (ii)). b) net relatedness index (NRI) and nearest taxon index (NTI) of all meta-sample groups in this meta-analysis.

### 3.4 Null Modelling

The NST and MST displayed in Figure 5 quantify the relative importance of stochasticity for each meta-sample group. Phylogenetic distances were calculated using Jaccard with and without abundances (Ruzicka approach). Both measures produced comparable results. NST values of >0.5 are more stochastic than deterministic. The highest NST values were in DWDS bulk water (0.71) followed closely by DWDS biofilm samples (0.59). Bulk water samples before and throughout treatment (up to disinfection) were also more stochastic than deterministic (0.53 and 0.58). Only two of the groups had more deterministic values: treated water (0.46) and biofilm in the water treatment works had the most deterministic value (0.43). MST values (modified ratio) are much lower, showing a similar pattern of values, except with untreated water, which has the highest (most stochastic) value (0.38).

**Figure 5:**
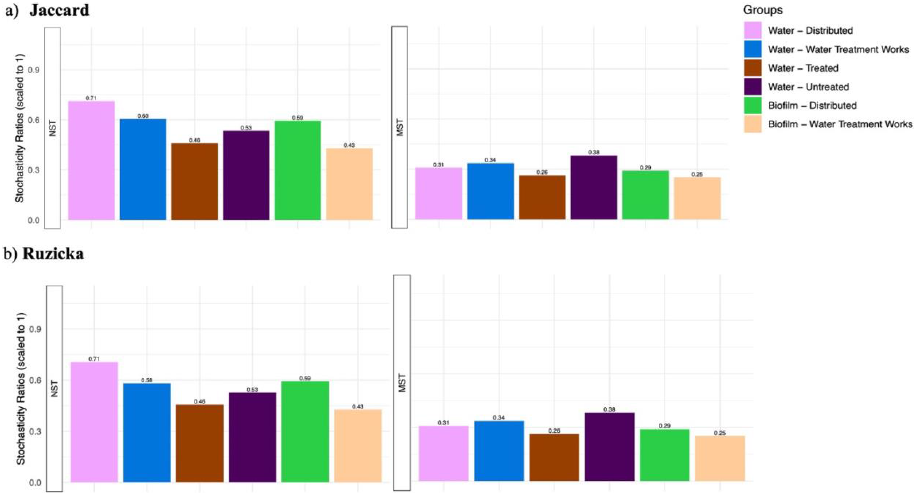
Normalised and Modified Stochasticity Ratios (NST/MST): **NST and MST** values for all meta-sample groups using a): Jaccard measures of phylogenetic distance and b) Ruzicka measures. Ruzicka is as Jaccard except that relative abundances are not considered.

### 3.5 Boolean Relationships

The results of the boolean analysis to identify individual relationships between ASVs in the dataset at 2 and 3-dimensional levels are displayed in Figure 6.

**Figure 6:**
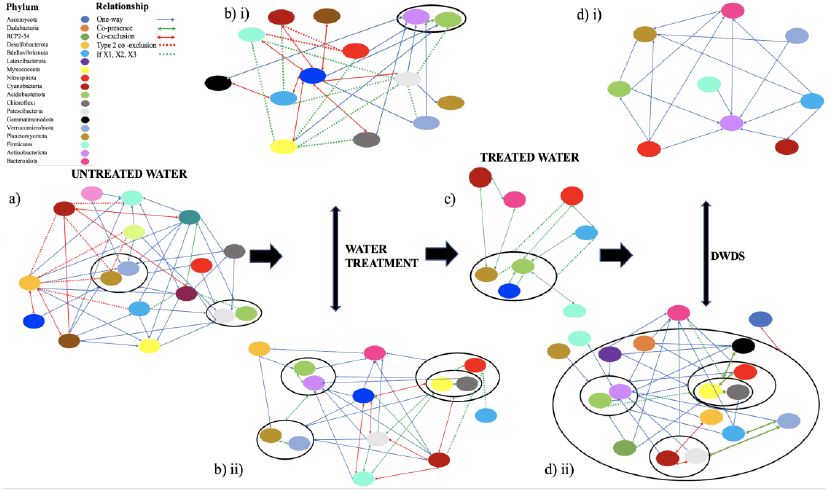
Phyla Relationships: Visual representation of individual Boolean relationships identified between Phyla for all meta-sample groups in this study, using methodology from Golovko et al. (2020). Minimal presence threshold set at 0.05% for presence-based relationships and maximum at 0.1% for absence ones. Groups are a) Untreated water; b) i) samples taken from bulk water throughout treatment; b) ii) biofilm from water treatment processes; c) disinfected water; d) i) water from distribution systems and d) ii) biofilm from pipes in distribution. 2-dimensional relationships detected included one-way relationships (blue arrow); co-presence (green arrow), and co-exclusion (red arrow); 3-dimensional relationships if ASV 1 is present, ASV 2 and 3 are also present (green dashed line) and type 2 co-exclusion (If ASV1 present then ASV2 and ASV3 are absent) (red dashed line). The black ellipses denote phyla that form similar relationships. NB *Bacteroidota* = *Bacteroidetes*.

The analysis identified 257 relationships between individual phyla across all stages of the treatment and distribution process, many of which were present across several groups. Most of the relationships found were one-way, although there were 21 three-dimensional relationships across all groups. These were predominantly found in untreated, water treatment and treated water groups. Treated water and bulk in water in DWDS had the least number of relationships, and no three-dimensional relationships were found in the DWDS group (11 and 17 respectively). Untreated water had 5 type 2 co-exclusionary relationships (if ASV1 then ASV2 and 3 are absent) detected. *Desulfobacterota* was involved in 3 of these types of relationships with *Bdellovibrionata* and *Cyanobacteria. Cyanobacteria* were also involved in several co-exclusionary relationships across the dataset and part of the only type 2 co-exclusionary relationship detected within the DWDS biofilm group (with *Desulfobacterota* and *Patescibacteriota*). Many of *Cyanobacteria*’s relationships were maintained across the groups, with both one-way and co-exclusion being most common, with one exception. *Cyanobacteria* were found to be in a co-presence relationship with *Bacteroidetes* and *Planctomycetes* in treated water, one of only 13 relationships detected for this group.

## 4. Discussion

### 4.1 Principles governing microbiomes in DWDS

Taxonomic profiles of the various meta-sample groups identified significant differences in abundances of genera throughout the source, treatment, and distribution of water. There were high degrees of species richness and alpha diversity of taxa in source waters that then reduced throughout treatment processes. This reduction is consistent with both the individual studies included in this analysis and several other pyrosequencing studies (Bautista-De los Santos et al., 2016; Pinto et al., 2014; Potgieter and Pinto, 2019). The reduction in LCBD in bulk water from untreated to treated water samples using Unifrac supports the reduction in alpha diversity and richness. A wide-scale study of 49 DWDS in China suggested reduced diversity and richness in tap compared to source waters (Han et al., 2020). The similarity of taxa increases throughout treatment and distribution sample groups, indicative of selective processes driven by filtration and chlorination, which only some organisms can survive (Hou et al., 2018; Lautenschlager et al., 2014; Vignola et al., 2018; Wan et al., 2019; Wang et al., 2014). However, LCBD values using Bray-Curtis dissimilarities increased across these groups, showing precisely the opposite pattern. It should be noted that Unifrac measures include information on phylogenetic relatedness which Bray Curtis does not. This highlights that beta diversity patterns throughout water treatment are incredibly complex and should be interpreted using several metrics.

Modelling of microbiomes now concentrates on the relative importance of random events on assembly, such as births, deaths, and environmental disturbance. This is compared to more traditional view that deterministic events such as selection drive communities (Sloan et al., 2006). As treatment and source choice may be significant in determining the organisms in the treated water, this is an important measure to consider. All meta-sample groups, except treated water and distributed biofilm, had more stochastic values suggesting a greater degree of randomness in microbiome assembly in these groups.

However, samples immediately after disinfection with chlorine had a more deterministic value. *Proteobacteria*, in particular: *Pseudomonas, Actinobacter*, and *Rheinheimera* have been demonstrated to dominate post disinfection, supporting the deterministic influence of treatment (Becerra-Castro et al., 2016). It also suggests that although filtration is important in defining taxa in DWDS, chlorine has a more strongly selective effect. This is supported by a study comparing two identical treatment systems treating the same source water, where chlorine and chloramine produced different bacterial communities (Potgieter and Pinto, 2019). In 2016, a meta-analysis of 14 pyrosequencing studies of water distribution systems compared the bacterial communities present under different disinfectant regimes, also confirming that the microbial communities in DWDS without a free chlorine residual are more diverse and abundant than those containing free chlorine.

*Mycobacterium* and *Pseudomonas* were significantly reduced by the presence of a free chlorine residual in that research (Bautista-de los Santos et al., 2016). In contrast, this meta-analysis found *Mycobacterium* was highly abundant in DWDS containing free chlorine and chloramine, but less so in those containing free chlorine residuals. *Pseudomonas* was only highly abundant in biofilm samples but absent in bulk water. It was more abundant in samples containing free chlorine, than chloramine. These differences may be due to sequencing technology, sample availability and sample site characteristics, and further highlight the complexity of interactions in the microbiomes of DWDS. These results do agree that *Pseudomonas* and *Mycobacterium* are important members of DWDS communities in general, however, and should be explored further to determine how these may be minimised, particularly considering their public health implications. This will be explored later in the discussion.

Bulk water in DWDS had the lowest LCBD using both metrics. The sharp reduction in LCBD in DWDS compared with at the outlet of the treatment process (treated water) suggests that although treatment is important in reducing diversity, time and space are important in shaping beta-diversity patterns in DWDS water. This is consistent with several previous studies on DWDS microbial communities, where temporal fluctuation had a significant impact on communities (McCoy and VanBriesen, 2012; Prest et al., 2014). There may also be potential diurnal cycles in bulk water, potentially due to flow patterns (Bautista-de los Santos et al., 2016). Many studies have also suggested that communities within DWDS are unique to that system, influenced by the source and treatment characteristics (Potgieter and Pinto, 2019; Wu et al., 2014). Although, the results presented here suggest that while the members of the community may be unique there is a general reduction in beta-diversity from treated to DWDS bulk water, indicating the efficacy of these processes.

Biofilm samples have a much higher LCBD than that of bulk water. These groups are also quite different in terms of types and abundance of taxa, a finding shared by individual studies (Douterelo et al., 2019, 2017; Liu et al., 2017b; Lührig et al., 2015). The core microbiome analysis for both these groups had only one organism in common, and the DWDS samples had lower abundances. This suggests that biofilm microbiomes contribute more to overall biodiversity within the pipe than the bulk water samples. This is important to consider, as the biofilm contains a quite different microbial profile than that of the bulk water, and sampling only bulk water may give a limited picture of the overall microbiome. This might lead to missed opportunities to detect pathogens. As aforementioned, *Pseudomonas* was the most abundant organism within biofilm but absent in bulk water. This is consistent with *Pseudomonas* as a primary coloniser of water biofilms (Doğruöz et al., 2009). Biofilm deposition is known to influence bulk water communities when loose deposits or biofilm are disturbed (Douterelo et al., 2019, 2017; Liu et al., 2017a). Biofilms can also significantly contribute to microbial loading in DWDS, with the composition affected by the presence of a chlorine residual (Fish and Boxall, 2018). These results were supported in this study by the high number of relationships evident between phyla in biofilm compared to bulk water (71:17 relationships identified). This, in addition to the elevated NRI and NTI values, suggest complex interactions define the assembly of biofilm communities.

A null modelling approach was also adopted to assess the importance of ecological stochasticity on the assembly of the microbiomes, using the NST and MST ratios (Ning et al., 2019). Assessing the relative importance of stochasticity has been applied to crude oil degrading marine bacterioplankton and anaerobic digestor communities using this approach and has been useful in understanding their assembly (Nikolova et al., 2021; Trego et al., 2021). The biofilm group from water treatment works had an NST value below 0.5, indicating than deterministic effects are more prominent in these samples, as they are close to null expectation. Although DWDS biofilm had more stochastic values, with higher NST values than the bulk water samples. These samples are more similar or dissimilar than the null expectation. The results from this dataset further supports the hypothesis that the effects of treatment and source reduce with distance and time from treatment (Han et al., 2020; Pinto et al., 2014; Potgieter et al., 2018). The strong selective pressures exerted by treatment processes therefore may be less important in DWDS.

The increased stochasticity in the DWDS group could be due to many environmental factors, including; genetic drift; the historical effects of treatment and source water on the assembly of the community including priority effects; the material and conditions within the pipe affecting dispersal; and flow conditions within the DWDS. This has been demonstrated by laboratory experiments using experimental pipe loops with the same influent water under different flow rates. The resulting biofilms contained some shared core taxa but with differences in their relative abundances, influenced by the flow conditions within each loop (Douterelo et al., 2017). Environmental impacts like priority effects, drift, dispersal and dispersal limitations may therefore be important factors in DWDS communities. These are important considerations in microbial assembly (Zhou and Ning, 2017). Long-term studies of discrete systems would be required to assess these impacts, at various temporal scales. Identification of significant one, two, and three-way relationships between individual Phyla in the meta-sample groups also demonstrates the complexity of DWDS microbiomes. 246 two-dimensional relationships were detected across all sample groups, the majority of these being one-way. There were also 21 three-dimensional relationships. These were particularly conserved across sample groups, with *Desulfobacterota*, *Cyanobacteria* and *Bdellovibrionata* forming co-exclusionary relationships in several groups. *Cyanobacteria* were also found to exclude *Bacteroidetes* in biofilm DWDS. *Bacteroidetes* is a phylum that contains pathogenic microbes, and while there are 6 orders within this phylum consisting of over 7000 species, the presence of these relationships suggests that further exploration of the abundance of these phyla is required (Thomas et al., 2011). It would be ideal to be able to assess these boolean relationships at a lower taxonomic resolution, but due to the significant increase in the number of relationships and reduced confidence values at lower levels, phylum provided the best resolution for this large data set. More targeted studies with a smaller sample size may be able to provide higher taxonomic resolution to explore important taxa relationships further.

### 4.2 Applying 16S rRNA Amplicon Sequencing for Water Utilities

As expected, the 25 most abundant taxa in the combined analysis did not contain any organisms traditionally used to indicate contamination, as these should be in low abundance in treated water. There were no coliforms identified in the core microbiome analysis or taxa profile for any meta-sample groups. Only one genus of *Enterobacteria* was detected in the most abundant taxa profiles from the individual datasets before collation (Supplementary information 3). *Citrobacter* - a member of the coliform group - was detected in around 10 samples, further suggesting that coliforms are in low abundance in water systems. It should be noted that analysis of untreated water samples in isolation failed to identify any highly abundant *Enterobacteriaceae*. Their low abundance in source waters may mean that 16S rRNA amplicon sequencing is not appropriate for the detection of traditional indicator organisms. These results also suggested indicators are not a part of the microbiome under normal operating conditions, although whether their detection is truly indicative of contamination was not assessed. Coliform bacteria are considered indicators of process performance, rather than faecal contamination (except *E. coli*) due to their prevalence in some environments and lack of correlation to other enteric pathogens (Ishii et al., 2006; Ishii and Sadowsky, 2008; Savichtcheva and Okabe, 2006). This study further suggests that their overall lack of abundance in untreated water makes them a poor indicator of process performance.

This analysis did reveal some organisms of concern as abundant in DWDS, although different organisms were of concern in different DWDS, consistent with the proposed system-specific nature of DWDS microbiomes (Roeselers et al., 2015). Of note, *Mycobacterium* was abundant in both chlorinated and chloraminated DWDS but was not prevalent in non-chlorinated DWDS samples. *Mycobacterium* is an emerging pathogen of concern for water utilities and dominates in some DWDS (Ashbolt, 2015; Zhu et al., 2019). *Nitrosomonas* and *Nitrospira* were also highly prevalent in the biofilm of chloraminated DWDS and an abundant member in several groups core microbiome analysis, supporting the results of individual studies highlighting their importance (Chan et al., 2019; Shaw et al., 2015). Monitoring the relative abundance of organisms like *Nitrosomonas* and *Nitrospira* using 16S rRNA amplicon sequencing may help utilities assess the risk of nitrification within chloraminated DWDS. Nitrite is an important regulatory parameter, but nitrification also indicates that disinfection levels are not sufficient to prevent microbial regrowth. Monitoring these groups of bacteria may help utilities intervene in DWDS before these issues become significant, through mains repair, flushing or replacement.

If water utilities can optimise processes to select for non-pathogenic microbes, this can reduce the risk of illness from drinking water, something suggested in several studies (Douterelo et al., 2019, 2017; Fish and Boxall, 2018). An overview of microbial communities’ dynamics, as ascertained by 16S rRNA amplicon analysis provides a holistic view of the response of water treatment and DWDS on microbiology. This understanding will aid risk management to maintain water quality and reduce the likelihood of harmful pathogens entering the system.

16S rRNA studies are becoming more popular and routine for the molecular analysis of water treatment and distribution. As demonstrated here, they provide extensive information revealing diverse communities that are influenced by the treatment process. However, translating this information into practice to inform and predict water quality is not always obvious to water utilities. However, taking a global meta-analysis view, this analysis highlighted several ways in which water utilities might employ 16S rRNA sequencing to improve drinking water quality. Considering whole microbial community dynamics from source to water, bulk and biofilm, this review has identified several organisms highly abundant throughout source and treatment, that can be potentially used to benchmark performance and monitor risk. As aforementioned, *Pseudomonas* and *Mycobacterium* were all abundant in DWDS, while *Legionella* was abundant in the source and treatment groups. Members of this genus are known pathogens of concern for drinking water quality. In particular, the higher abundance of the genus *Legionella* in source waters and treatment in this analysis may make it a good indicator of treatment performance, especially as other studies have detected *Legionella* in treated water samples (Ashbolt, 2015; Hou et al., 2018; Vignola et al., 2018). *Legionella* is an emerging pathogen of concern to the water industry, and in the UK may include this in future water quality regulations. The *Legionella* genus contains 71 species, only a few of which are pathogenic, with *Legionella pneumophilae* being of most concern (Doleans et al., 2004; Garrity et al., 2005). In this meta-analysis, 67 different sequences were assigned to *Legionella* species, including *L. pneumophilae*, *L. longbeachae and L. bozemanii*. These species account for 96.3% of cases of legionellosis worldwide (Doleans et al., 2004). Therefore, amplicon sequencing can aid water utilities to assess the risk of legionellosis from water systems, although the viability of the organisms detected must also be considered using an alternative method.

Flow Cytometry (FCM) may provide the appropriate information to compliment 16S rRNA sequencing. Using FCM with the intercalating dyes SYBr Green and propidium iodide to stain genetic material *in situ* within a sample gives a quantitative measure of the intact cells within the microbiome of DWDS. This is due to the ability of SYBr green to permeate intact cell membranes, whereas propidium iodide cannot. Dead cells, therefore, appear red, where intact cells fluoresce green, allowing each to be distinguished. This approach has been extensively explored in studies assessing water treatment cell removal, DWDS regrowth and seasonal changes within microbiomes (Besmer and Hammes, 2016; Hammes et al., 2010; Hassard et al., 2019; Prest et al., 2016). FCM can also provide more information than just the count of cells within a sample, using the relative fluorescence and a statistical binning process, cells can be grouped into populations which can then be tracked (Favere et al., 2020; Props et al., 2016). A quantitative measure like the ratio of intact cells to the total count of cells within a sample could be used by utilities to quantify the viability of the organisms identified using a 16S rRNA amplicon sequencing, enhancing the benefits of both analyses.

Measures of species richness and abundance such as alpha diversity and LCBD at treatment and distribution stages are useful to water utilities when comparing DWDS performance. Although monitoring the relative abundance of specific taxa in a single DWDS may not be able to detect a risk to public health directly, an understanding of these values across different DWDS allow water utilities to assess the impacts of source, treatment and distribution conditions on water quality and make more informed choices on asset investment. Understanding the relative impacts of ecological stochasticity in DWDS microbiomes is also a useful exercise for water utilities. Higher stochastic values in bulk water and biofilm of DWDS samples in this analysis suggest that random events are more important than treatment processes or other prior deterministic events in determining the bacterial communities. In contrast, biofilm communities in the water treatment works and bulk treated water groups are more deterministic, affected by the abiotic conditions (e.g., chlorine, pH, pipe material) and prior treatment processes. Managing biofilm and ensuring treatment processes remove microbes of concern is where water utilities can most effectively minimise risk to public health.

This discussion has focussed primarily on the results from DWDS, as this provided the largest proportion of the available data. There is a need to sample further at source and throughout treatment processes to explore the ecological rules and relationships between taxa at these stages and their subsequent impact on assembly of microbiomes in the DWDS. This is a particularly important question for water utilities so that they can improve treatment and distribution processes.

## 5. Conclusions

There are copious quantities of data from amplicon sequencing studies in DWDS. Although many of these studies provide only a descriptive understanding of the microbiomes. As a result, this information has yet to be used to predict and direct microbial water quality. There has been a reluctance to adopt sequencing technology among water utilities, as the benefits are not immediately clear. Using a meta-analysis of 16S rRNA amplicons, we have shown that while treatment and distribution of water significantly reduce the diversity and abundance of taxa present in the source, the subsequent assembly of microbiomes in drinking water is a stochastic process, particularly in the DWDS. This suggests that the effects of source and treatment diminish with distance and time. Only the assembly of microbiomes at the point of chlorination is more deterministic, due to selection pressures on organisms that cannot survive oxidation. Although 16S rRNA amplicon sequencing cannot satisfy current water quality regulation, it can assess the risk from emerging pathogens such as *Legionella* or *Mycobacterium* or biofilm colonisers like *Pseudomonas* - which may be in high abundance in DWDS. Amplicon sequencing can also track significant changes in the microbiome, which may be associated with contamination or changes in process performance. These are benefits traditional culture tests cannot provide. However, further work is required to standardise sequencing and data analysis methods for 16S rRNA amplicon sequencing methods to enable the water industry to adopt them as standard practice.

## Acknowledgements

Funding: CT. Scottish Water Industry Funded PhD studentship; by EPSRC award EP/V030515/1, a Royal Academy of Engineering-Scottish Water Research Chair (RCSRF171864) awarded to CJS, and UZI is supported by NERC, UK, NE/L011956/1.

---

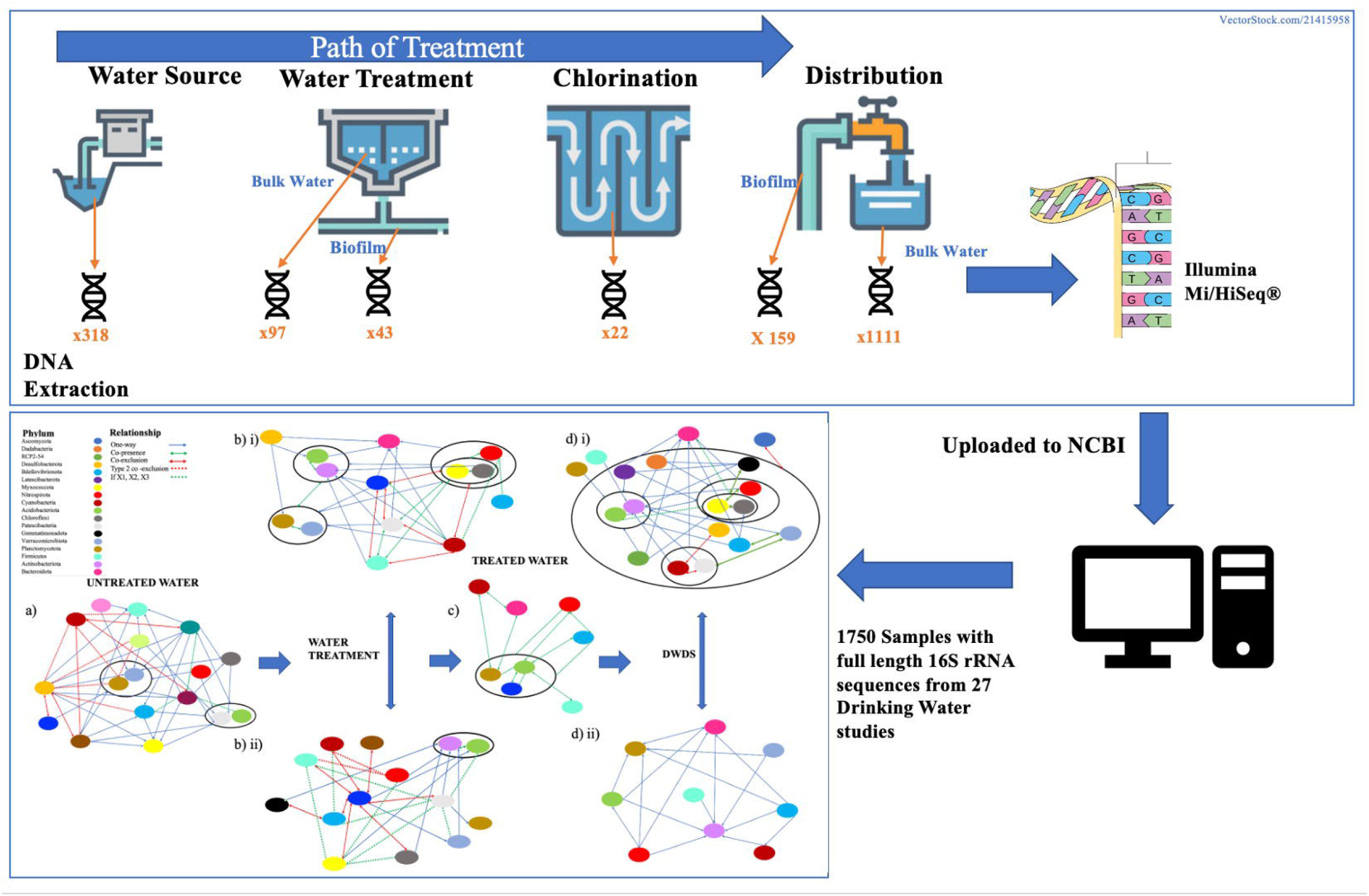

## References

Ahmed, W., Staley, C., Sadowsky, M.J., Gyawali, P., Sidhu, J.P.S., Palmer, A., Beale, D.J., Toze, S., 2015. Toolbox Approaches Using Molecular Markers and 16S rRNA Gene Amplicon Data Sets for Identification of Fecal Pollution in Surface Water. Appl. Environ. Microbiol. 81, 7067–7077. https://doi.org/10.1128/AEM.02032-15

Ashbolt, N.J., 2015. Microbial Contamination of Drinking Water and Human Health from Community Water Systems. Curr. Environ. Heal. reports. https://doi.org/10.1007/s40572-014-0037-5

Bautista-de los Santos, Q.M., Schroeder, J.L., Blakemore, O., Moses, J., Haffey, M., Sloan, W., Pinto, A.J., 2016. The impact of sampling, PCR, and sequencing replication on discerning changes in drinking water bacterial community over diurnal time-scales. Water Res. 90, 216–224. https://doi.org/10.1016/j.watres.2015.12.010

Bautista-De los Santos, Q.M., Schroeder, J.L., Sevillano-Rivera, M.C., Sungthong, R., Ijaz, U.Z., Sloan, W.T., Pinto, A.J., 2016. Emerging investigators series: microbial communities in full-scale drinking water distribution systems – a meta-analysis. Environ. Sci. Water Res. Technol. 2, 631–644. https://doi.org/10.1039/C6EW00030D

Becerra-Castro, C., Macedo, G., Silva, A.M.T., Manaia, C.M., Nunes, O.C., 2016. Proteobacteria become predominant during regrowth after water disinfection. Sci. Total Environ. 573, 313–323. https://doi.org/10.1016/j.scitotenv.2016.08.054

Besmer, M.D., Hammes, F., 2016. Short-term microbial dynamics in a drinking water plant treating groundwater with occasional high microbial loads. Water Res. 107, 11–18. https://doi.org/10.1016/j.watres.2016.10.041

Cabral, J.P.S., 2010. Water microbiology. Bacterial pathogens and water. Int. J. Environ. Res. Public Health 7, 3657–3703. https://doi.org/10.3390/ijerph7103657

Chan, S., Pullerits, K., Keucken, A., Persson, K.M., Paul, C.J., Rådström, P., 2019. Bacterial release from pipe biofilm in a full-scale drinking water distribution system. npj Biofilms Microbiomes 5, 9. https://doi.org/10.1038/s41522-019-0082-9

D’Amore, R., Ijaz, U.Z., Schirmer, M., Kenny, J.G., Gregory, R., Darby, A.C., Shakya, M., Podar, M., Quince, C., Hall, N., 2016. A comprehensive benchmarking study of protocols and sequencing platforms for 16S rRNA community profiling. BMC Genomics 17, 1–20. https://doi.org/10.1186/s12864-015-2194-9

De Vera, G.A., Gerrity, D., Stoker, M., Frehner, W., Wert, E.C., 2018. Impact of upstream chlorination on filter performance and microbial community structure of GAC and anthracite biofilters. Environ. Sci. Water Res. Technol. 4, 1133–1144. https://doi.org/10.1039/c8ew00115d

Doğruöz, N., Göksay, D., Ilhan-Sungur, E., Cotuk, A., 2009. Pioneer colonizer microorganisms in biofilm formation on galvanized steel in a simulated recirculating cooling-water system. J. Basic Microbiol. 49. https://doi.org/10.1002/jobm.200800250

Doleans, A., Aurell, H., Reyrolle, M., Lina, G., Freney, J., Vandenesch, F., Etienne, J., Jarraud, S., 2004. Clinical and Environmental Distributions of Legionella Strains in France Are Different. J. Clin. Microbiol. 42, 458. https://doi.org/10.1128/JCM.42.1.458-460.2004

Douterelo, I., Jackson, M., Solomon, C., Boxall, J., 2017. Spatial and temporal analogies in microbial communities in natural drinking water biofilms. Sci. Total Environ. 581–582, 277–288. https://doi.org/10.1016/j.scitotenv.2016.12.118

Douterelo, I., Sharpe, R.L., Husband, S., Fish, K.E., Boxall, J.B., 2019. Understanding microbial ecology to improve management of drinking water distribution systems. Wiley Interdiscip. Rev. Water 6, e01325. https://doi.org/10.1002/wat2.1325

DWI, 2020. Public Drinking Water Quality 2020 [WWW Document]. Annu. Rep. URL https://www.dwi.gov.uk/what-we-do/annual-report/drinking-water-2020/%0Ahttps://www.dwi.gov.uk/ (accessed 6.17.21).

Favere, J., Buysschaert, B., Boon, N., De Gusseme, B., 2020. Online microbial fingerprinting for quality management of drinking water: Full-scale event detection. Water Res. 170. https://doi.org/10.1016/j.watres.2019.115353

Fish, K.E., Boxall, J.B., 2018. Biofilm Microbiome (Re)Growth Dynamics in Drinking Water Distribution Systems Are Impacted by Chlorine Concentration. Front. Microbiol. 9, 2519. https://doi.org/10.3389/fmicb.2018.02519

Fox, E.J., Reid-Bayliss, K.S., 2014. Accuracy of Next Generation Sequencing Platforms. J. Next Gener. Seq. Appl. 01. https://doi.org/10.4172/2469-9853.1000106

Fricker, C.R., Eldred, B.J., 2009. Identification of coliform genera recovered from water using different technologies. Lett. Appl. Microbiol. 49, 685–688. https://doi.org/10.1111/j.1472-765X.2009.02726.x

Garrity, G.M., Bell, J.A., Lilburn, T., 2005. Legionellales ord. nov., in: Brenner, D.J., Krieg, N.R., Staley, J.T., Garrity, G.M., Boone, D.R., De Vos, P., Goodfellow, M., Rainey, F.A., Schleifer, K.-H. (Eds.), Bergey’s Manual^®^ of Systematic Bacteriology. Springer US, Boston, MA, pp. 210–247. https://doi.org/10.1007/0-387-28022-7_6

Gerrity, D., Arnold, M., Dickenson, E., Moser, D., Sackett, J.D., Wert, E.C., 2018. Microbial community characterization of ozone-biofiltration systems in drinking water and potable reuse applications. Water Res. 135, 207–219. https://doi.org/10.1016/j.watres.2018.02.023

Ghaju Shrestha, R., Tanaka, Y., Malla, B., Bhandari, D., Tandukar, S., Inoue, D., Sei, K., Sherchand, J.B., Haramoto, E., 2017. Next-generation sequencing identification of pathogenic bacterial genes and their relationship with fecal indicator bacteria in different water sources in the Kathmandu Valley, Nepal. Sci. Total Environ. 601–602, 278–284. https://doi.org/10.1016/j.scitotenv.2017.05.105

Gülay, A., Musovic, S., Albrechtsen, H.J., Al-Soud, W.A., Sørensen, S.J., Smets, B.F., 2016. Ecological patterns, diversity and core taxa of microbial communities in groundwater-fed rapid gravity filters. ISME J. 10, 2209–2222. https://doi.org/10.1038/ismej.2016.16

Hahn, M.W., Koll, U., Schmidt, J., 2019. Isolation and Cultivation of Bacteria, in: The Structure and Function of Aquatic Microbial Communities. Springer, Cham, pp. 313–351. https://doi.org/10.1007/978-3-030-16775-2_10

Hammes, F., Goldschmidt, F., Vital, M., Wang, Y., Egli, T., 2010. Measurement and interpretation of microbial adenosine tri-phosphate (ATP) in aquatic environments. Water Res. 44, 3915–3923. https://doi.org/10.1016/j.watres.2010.04.015

Han, Z., An, W., Yang, M., Zhang, Y., 2020. Assessing the impact of source water on tap water bacterial communities in 46 drinking water supply systems in China. Water Res. 172, 115469. https://doi.org/10.1016/j.watres.2020.115469

Hassard, F., Whitton, R., Jefferson, B., Jarvis, P., 2019. Understanding the Use of Flow Cytometry for Monitoring of Drinking Water, The UK Drinking Water Inspectorate.

Hou, L., Zhou, Q., Wu, Q., Gu, Q., Sun, M., Zhang, J., 2018. Spatiotemporal changes in bacterial community and microbial activity in a full-scale drinking water treatment plant. Sci. Total Environ. 625, 449–459. https://doi.org/10.1016/j.scitotenv.2017.12.301

Ishii, S., Ksoll, W.B., Hicks, R.E., Sadowsky, M.J., 2006. Presence and growth of naturalized Escherichia coli in temperate soils from lake superior watersheds. Appl. Environ. Microbiol. 72, 612–621. https://doi.org/10.1128/AEM.72.1.612-621.2006

Ishii, S., Sadowsky, M.J., 2008. Escherichia coli in the Environment: Implications for Water Quality and Human Health. microbes and genomes 23, 101–108. https://doi.org/10.1264/jsme2.23.101

Jalava, K., Rintala, H., Ollgren, J., Maunula, L., Gomez-Alvarez, V., Revez, J., Palander, M., Antikainen, J., Kauppinen, A., Räsänen, P., Siponen, S., Nyholm, O., Kyyhkynen, A., Hakkarainen, S., Merentie, J., Pärnänen, M., Loginov, R., Ryu, H., Kuusi, M., Siitonen, A., Miettinen, I., Santo Domingo, J.W., Hänninen, M.L., Pitkänen, T., 2014. Novel microbiological and spatial statistical methods to improve strength of epidemiological evidence in a community-wide waterborne outbreak. PLoS One 9, 104713. https://doi.org/10.1371/journal.pone.0104713

Keating, C., Trego, A.C., Sloan, W., O’Flaherty, V., Ijaz, U.Z., 2020. Circular Economy of Anaerobic Biofilm Microbiomes: A Meta-Analysis Framework for Re-exploration of Amplicon Sequencing Data. bioRxiv 2020.12.23.424166. https://doi.org/10.1101/2020.12.23.424166

Kuusi, M., Nuorti, J.P., Hänninen, M.L., Koskela, M., Jussila, V., Kela, E., Miettinen, I., Ruutu, P., 2005. A large outbreak of campylobacteriosis associated with a municipal water supply in Finland. Epidemiol. Infect. 133, 593–601. https://doi.org/10.1017/S0950268805003808

Lahti, L., Sudarshan, S., et al., 2017. Introduction to the microbiome R package [WWW Document]. Bioconductor. URL https://microbiome.github.io/tutorials/ (accessed 4.7.21).

Lautenschlager, K., Hwang, C., Ling, F., Liu, W.T., Boon, N., Köster, O., Egli, T., Hammes, F., 2014. Abundance and composition of indigenous bacterial communities in a multi-step biofiltration-based drinking water treatment plant. Water Res. 62, 40–52. https://doi.org/10.1016/j.watres.2014.05.035

Leclerc, H., Mossel, D.A.A., Edberg, S.C., Struijk, C.B., 2001. Advances in the Bacteriology of the Coliform Group: Their Suitability as Markers of Microbial Water Safety. Annu. Rev. Microbiol. 55, 201–234. https://doi.org/10.1146/annurev.micro.55.1.201

Legendre, P., De Cáceres, M., 2013. Beta diversity as the variance of community data: Dissimilarity coefficients and partitioning. Ecol. Lett. 16, 951–963. https://doi.org/10.1111/ele.12141

Lin, W., Yu, Z., Zhang, H., Thompson, I.P., 2014. Diversity and dynamics of microbial communities at each step of treatment plant for potable water generation. Water Res. 52, 218–230. https://doi.org/10.1016/j.watres.2013.10.071

Liu, G., Tao, Y., Zhang, Y., Lut, M., Knibbe, W.J., van der Wielen, P., Liu, W., Medema, G., van der Meer, W., 2017a. Hotspots for selected metal elements and microbes accumulation and the corresponding water quality deterioration potential in an unchlorinated drinking water distribution system. Water Res. 124, 435–445. https://doi.org/10.1016/j.watres.2017.08.002

Liu, G., Zhang, Y., Knibbe, W.J., Feng, C., Liu, W., Medema, G., van der Meer, W., 2017b. Potential impacts of changing supply-water quality on drinking water distribution: A review. Water Res. https://doi.org/10.1016/j.watres.2017.03.031

Lührig, K., Canbäck, B., Paul, C.J., Johansson, T., Persson, K.M., Rådström, P., 2015. Bacterial community analysis of drinking water biofilms in southern Sweden. Microbes Environ. 30, 99–107. https://doi.org/10.1264/jsme2.ME14123

McCoy, S.T., VanBriesen, J.M., 2012. Temporal Variability of Bacterial Diversity in a Chlorinated Drinking Water Distribution System. J. Environ. Eng. 138, 786–795. https://doi.org/10.1061/(asce)ee.1943-7870.0000539

Nearing, J.T., Douglas, G.M., Comeau, A.M., Langille, M.G.I., 2018. Denoising the Denoisers: An independent evaluation of microbiome sequence error-correction approaches. PeerJ 2018. https://doi.org/10.7717/peerj.5364

Nikolova, C., Ijaz, U.Z., Gutierrez, T., 2021. Exploration of marine bacterioplankton community assembly mechanisms during chemical dispersant and surfactant-assisted oil biodegradation. Ecol. Evol. 11, 13862–13874. https://doi.org/10.1002/ece3.8091

Ning, D., Deng, Y., Tiedje, J.M., Zhou, J., 2019. A general framework for quantitatively assessing ecological stochasticity. Proc. Natl. Acad. Sci. U. S. A. 116, 16892–16898. https://doi.org/10.1073/pnas.1904623116

Payment, P., Locas, A., 2011. Pathogens in Water: Value and Limits of Correlation with Microbial Indicators. Ground Water 49, 4–11. https://doi.org/10.1111/j.1745-6584.2010.00710.x

Pinto, A.J., Schroeder, J., Lunn, M., Sloan, W., Raskin, L., 2014. Spatial-Temporal Survey and Occupancy-Abundance Modeling To Predict Bacterial Community Dynamics in the Drinking Water Microbiome. MBio 5. https://doi.org/10.1128/mBio.01135-14

Pinto, A.J., Xi, C., Raskin, L., 2012. Bacterial Community Structure in the Drinking Water Microbiome Is Governed by Filtration Processes. Environ. Sci. Technol. 46, 8851–8859. https://doi.org/10.1021/es302042t

Potgieter, S., Pinto, A., Sigudu, M., du Preez, H., Ncube, E., Venter, S., 2018. Long-term spatial and temporal microbial community dynamics in a large-scale drinking water distribution system with multiple disinfectant regimes. Water Res. 139, 406–419. https://doi.org/10.1016/j.watres.2018.03.077

Potgieter, S.C., Pinto, A.J., 2019. Reproducible microbial community dynamics of two drinking water systems treating similar source waters. bioRxiv Microbiol. 678920. https://doi.org/10.1101/678920

Prest, E.I., El-Chakhtoura, J., Hammes, F., Saikaly, P.E., van Loosdrecht, M.C.M., Vrouwenvelder, J.S., 2014. Combining flow cytometry and 16S rRNA gene pyrosequencing: A promising approach for drinking water monitoring and characterization. Water Res. 63, 179–189. https://doi.org/10.1016/j.watres.2014.06.020

Prest, E.I., Hammes, F., Kötzsch, S., Van Loosdrecht, M.C.M., Vrouwenvelder, J.S., 2016. A systematic approach for the assessment of bacterial growth-controlling factors linked to biological stability of drinking water in distribution systems. Water Sci. Technol. Water Supply 16, 865–880. https://doi.org/10.2166/ws.2016.001

Props, R., Monsieurs, P., Mysara, M., Clement, L., Boon, N., 2016. Measuring the biodiversity of microbial communities by flow cytometry. Methods Ecol. Evol. 7, 1376–1385. https://doi.org/10.1111/2041-210X.12607

Rappé, M.S., Giovannoni, S.J., 2003. The Uncultured Microbial Majority. Annu. Rev. Microbiol. https://doi.org/10.1146/annurev.micro.57.030502.090759

Roeselers, G., Coolen, J., van der Wielen, P.W.J.J., Jaspers, M.C., Atsma, A., de Graaf, B., Schuren, F., 2015. Microbial biogeography of drinking water: Patterns in phylogenetic diversity across space and time. Environ. Microbiol. 17, 2505–2514. https://doi.org/10.1111/1462-2920.12739

Savichtcheva, O., Okabe, S., 2006. Alternative indicators of fecal pollution: Relations with pathogens and conventional indicators, current methodologies for direct pathogen monitoring and future application perspectives. Water Res. https://doi.org/10.1016/j.watres.2006.04.040

Saxena, G., Bharagava, R.N., Kaithwas, G., Raj, A., 2015. Microbial indicators, pathogens and methods for their monitoring in water environment. J. Water Health 13, 319–339. https://doi.org/10.2166/wh.2014.275

Schirmer, M., Ijaz, U.Z., D’Amore, R., Hall, N., Sloan, W.T., Quince, C., 2015. Insight into biases and sequencing errors for amplicon sequencing with the Illumina MiSeq platform. Nucleic Acids Res. 43. https://doi.org/10.1093/nar/gku1341

Shaw, J.L.A.A., Monis, P., Weyrich, L.S., Sawade, E., Drikas, M., Cooper, A.J., 2015. Using Amplicon Sequencing To Characterize and Monitor Bacterial Diversity in Drinking Water Distribution Systems. Appl. Environ. Microbiol. 81, 6463–6473. https://doi.org/10.1128/AEM.01297-15

Sloan, W.T., Lunn, M., Woodcock, S., Head, I.M., Nee, S., Curtis, T.P., 2006. Quantifying the roles of immigration and chance in shaping prokaryote community structure. Environ. Microbiol. 8, 732–740. https://doi.org/10.1111/j.1462-2920.2005.00956.x

Thomas, F., Hehemann, J.H., Rebuffet, E., Czjzek, M., Michel, G., 2011. Environmental and gut Bacteroidetes: The food connection. Front. Microbiol. 2, 93. https://doi.org/10.3389/FMICB.2011.00093/BIBTEX

Trego, A.C., McAteer, P.G., Nzeteu, C., Mahony, T., Abram, F., Ijaz, U.Z., O’Flaherty, V., 2021. Combined Stochastic and Deterministic Processes Drive Community Assembly of Anaerobic Microbiomes During Granule Flotation. Front. Microbiol. 12, 1165. https://doi.org/10.3389/FMICB.2021.666584/BIBTEX

Uyaguari-Diaz, M.I., Croxen, M.A., Cronin, K., Luo, Z., Isaac-Renton, J., Prystajecky, N.A., Tang, P., 2019. Microbial community dynamics of surface water in British Columbia, Canada. bioRxiv 719146. https://doi.org/10.1101/719146

Vierheilig, J., Savio, D., Farnleitner, A.H., Reischer, G.H., Ley, R.E., Mach, R.L., Farnleitner, A.H., Reischer, G.H., 2015. Potential applications of next generation DNA sequencing of 16S rRNA gene amplicons in microbial water quality monitoring. Water Sci. Technol. 72, 1962–1972. https://doi.org/10.2166/wst.2015.407

Vignola, M., Werner, D., Wade, M.J., Meynet, P., Davenport, R.J., 2018. Medium shapes the microbial community of water filters with implications for effluent quality. Water Res. 129, 499–508. https://doi.org/10.1016/j.watres.2017.09.042

Wan, K., Zhang, M., Ye, C., Lin, W., Guo, L., Chen, S., Yu, X., 2019. Organic carbon: An overlooked factor that determines the antibiotic resistome in drinking water sand filter biofilm. Environ. Int. 125, 117–124. https://doi.org/10.1016/j.envint.2019.01.054

Wang, H., Masters, S., Edwards, M.A., Falkinham, J.O., Pruden, A., 2014. Effect of Disinfectant, Water Age, and Pipe Materials on Bacterial and Eukaryotic Community Structure in Drinking Water Biofilm. Environ. Sci. Technol. 48, 1426–1435. https://doi.org/10.1021/es402636u

Wang, H., Pryor, M.A., Edwards, M.A., Falkinham, J.O., Pruden, A., 2013. Effect of GAC pre-treatment and disinfectant on microbial community structure and opportunistic pathogen occurrence. Water Res. 47, 5760–5772. https://doi.org/10.1016/j.watres.2013.06.052

Wolf-Baca, M., Piekarska, K., 2020. Biodiversity of organisms inhabiting the water supply network of Wroclaw. Detection of pathogenic organisms constituting a threat for drinking water recipients. Sci. Total Environ. 715, 136732. https://doi.org/10.1016/j.scitotenv.2020.136732

Wu, H., Zhang, J., Mi, Z., Xie, S., Chen, C., Zhang, X., 2014. Biofilm bacterial communities in urban drinking water distribution systems transporting waters with different purification strategies. Appl. Microbiol. Biotechnol. 99, 1947–1955. https://doi.org/10.1007/s00253-014-6095-7

Zhou, J., Ning, D., 2017. Stochastic Community Assembly: Does It Matter in Microbial Ecology? Microbiol. Mol. Biol. Rev. 81, 1–32. https://doi.org/10.1128/MMBR.00002-17

Zhu, J., Liu, R., Cao, N., Yu, J., Liu, X., Yu, Z., 2019. Mycobacterial metabolic characteristics in a water meter biofilm revealed by metagenomics and metatranscriptomics. Water Res. 153, 315–323. https://doi.org/10.1016/j.watres.2019.01.032

