## Supplementary Information 1 for "Microbiomes in drinking water treatment and distribution: a meta-analysis from source to tap"

Table 1: List of the research papers whose data was used in this meta-analysis

|  | **Authors** | **Title** | **Year** | **Journal/Repository** | **DOI** | **Bioproject Number** |
| --- | --- | --- | --- | --- | --- | --- |
| 1 | Hou L., Zhou Q., Wu Q., Gu Q., Sun M., Zhang J. | Spatiotemporal changes in bacterial community and microbial activity in a full-scale drinking water treatment plant | 2018 | Science of the Total Environment | [10.1016/j.scitotenv.2017.12.301](https://doi.org/10.1016/j.scitotenv.2017.12.301) | PRJNA399213 |
| 2 | Wan K., Zhang M., Ye C., Lin W., Guo L., Chen S., Yu X. | Organic carbon: An overlooked factor that determines the antibiotic resistome in drinking water sand filter biofilm | 2019 | Environment International | [10.1016/j.envint.2019.01.054](https://doi.org/10.1016/j.envint.2019.01.054) | PRJNA451227 |
| 3 | Liu G., Tao Y., Zhang Y., Lut M., Knibbe W.-J., van der Wielen P., Liu W., Medema G., van der Meer W. | Hotspots for selected metal elements and microbes accumulation and the corresponding water quality deterioration potential in an unchlorinated drinking water distribution system | 2017 | Water Research | [10.1016/j.watres.2017.08.002](https://doi.org/10.1016/j.watres.2017.08.002) | PRJNA393048 |
| 4 | Liu G., Zhang Y., van der Mark E., Magic-Knezev A., Pinto A., van den Bogert B., Liu W., van der Meer W., Medema G. | Assessing the origin of bacteria in tap water and distribution system in an unchlorinated drinking water system by SourceTracker using microbial community fingerprints | 2018 | Water Research | [10.1016/j.watres.2018.03.043](https://doi.org/10.1016/j.watres.2018.03.043) | PRJNA397332 |
| 5 | Douterelo, I., Jackson, M., Solomon, C. & Boxall, J. | Spatial and temporal analogies in microbial communities in natural drinking water biofilm | 2017 | Science of the Total Environment | [10.1016/j.scitotenv.2016.12.118.](https://doi.org/10.1016/j.scitotenv.2016.12.118.) | PRJNA347544 |
| 6 | Fish K, and Boxall, J | Biofilm Microbiome (Re)Growth Dynamics in Drinking Water Distribution Systems Are Impacted by Chlorine Concentration | 2018 | Frontiers in Microbiology | [10.3389/fmicb.2018.02519](https://doi.org/10.3389/fmicb.2018.02519) | PRJNA479666 |
| 7 | Waak, M. B., Hozalski, R. M., Hallé, C. & LaPara, T. M. | Comparison of the microbiomes of two drinking water distribution systems—with and without residual chloramine disinfection | 2019 | Microbiome | <10.1186/s40168-019-0707-5> | PRJNA433427 |
| 8 | Perrin, Y., Bouchon, D., Delafont, V., Moulin, L. & Héchard, Y. | Microbiome of drinking water: A full-scale spatio-temporal study to monitor water quality in the Paris distribution system | 2018 | Water Research | [10.1016/j.watres.2018.11.013](https://doi.org/10.1016/j.watres.2018.11.013) | PRJEB24989 |
| 9 | Jasser I., Kostrzewska-Szlakowska I., Kwiatowski J., Navruzshoev D., Suska-Malawska M., Khomutovska N. | Morphological and Molecular Diversity of Benthic Cyanobacteria Communities Versus Environmental Conditions in Shallow, High Mountain Water Bodies in Eastern Pamir Mountains (Tajikistan) | 2019 | Polish Journal of Ecology | [10.3161/15052249PJE2019.67.4.002](https://doi.org/10.3161/15052249PJE2019.67.4.002) | PRJNA486727 |
| 10 | Chan, S., Pullerits, K., Keucken, A., Alexander P., K.M., Paul, C.J., Rådström, P. | Bacterial release from pipe biofilm in a full-scale drinking water distribution system | 2019 | NPJ Biofilms and Microbiomes | [10.1038/s41522-019-0082-9](https://doi.org/10.1038/s41522-019-0082-9) | PRJNA494637 |
| 11 | Wolf-Baca M., Piekarska K. | Biodiversity of organisms inhabiting the water supply network of Wroclaw. Detection of pathogenic organisms constituting a threat for drinking water recipients | 2020 | Science of the Total Environment | [10.1016/j.scitotenv.2020.136732](https://doi.org/10.1016/j.scitotenv.2020.136732) | PRJUOW1 |
| 12 | Bisht, G., Sourirajan, A., Baumler, D. J. & Dev, K. | 16S rRNA Gene Amplicon Data Set-Based Bacterial Diversity in a Water-Soil Sample from Pangong Tso Lake, a High-Altitude Grassland Lake of the Northwest Himalayas | 2018 | American Society for Microbiology | [10.1128/MRA.01192-18](https://doi.org/10.1128/MRA.01192-18) | PRJNA486366 |
| 13 | Zhu J., Liu R., Cao N., Yu J., Liu X., Yu Z. | Mycobacterial metabolic characteristics in a water meter biofilm revealed by metagenomics and metatranscriptomics | 2019 | Water Research | [10.1016/j.watres.2019.01.032](https://doi.org/10.1016/j.watres.2019.01.032) | PRJNA487710 |
| 14 | Ji, P., Parks, J., Edwards, M. A. & Pruden, A. | Impact of Water Chemistry, Pipe Material and Stagnation on the Building Plumbing Microbiome | 2015 | PLoS One | [10.1371/journal.pone.0141087](https://doi.org/10.1371/journal.pone.0141087) | QIITA 10251 |
| 15 | Roeselers, G., Coolen, J., van der Wielen, P.W.J.J., Jaspers, M.C., Atsma, A., de Graaf, B., Schuren, F. | Microbial biogeography of drinking water: patterns in phylogenetic diversity across space and time | 2015 | Environmental Microbiology | [10.1111/1462-2920.12739](https://doi.org/10.1111/1462-2920.12739) | PRJEB7435 |
| 16 | Uyaguari-Diaz, M. I., Croxen, M.A., Cronin, K., Luo, Z., Isaac-Renton, J., Prystajecky, N.A., Tang, P. | Microbial community dynamics of surface water in British Columbia, Canada | 2019 | bioRxiv | [10.1101/719146](https://doi.org/10.1101/719146) | PRJNA287840 |
| 17 | Ghaju Shrestha R., Tanaka Y., Malla B., Bhandari D., Tandukar S., Inoue D., Sei K., Sherchand J.B., Haramoto E. | Next-generation sequencing identification of pathogenic bacterial genes and their relationship with fecal indicator bacteria in different water sources in the Kathmandu Valley, Nepal | 2017 | Science of the Total Environment | [10.1016/j.scitotenv.2017.05.105](https://doi.org/10.1016/j.scitotenv.2017.05.105) | PRJDB5406 |
| 18 | Bautista-de los Santos Q.M., Schroeder J.L., Blakemore O., Moses J., Haffey M., Sloan W., Pinto A.J. | The impact of sampling, PCR, and sequencing replication on discerning changes in drinking water bacterial community over diurnal time-scales | 2016 | Water Research | [10.1016/j.watres.2015.12.010](https://doi.org/10.1016/j.watres.2015.12.010) | PRJNA283789 |
| 19 | Potgieter S., Pinto A., Sigudu M., du Preez H., Ncube E., Venter S. | Long-term spatial and temporal microbial community dynamics in a large-scale drinking water distribution system with multiple disinfectant regimes | 2018 | Water Research | [10.1016/j.watres.2018.03.077](https://doi.org/10.1016/j.watres.2018.03.077) | PRJNA445682 |
| 20 | Jalava, K., Rintala, H., Ollgren, J., Maunula, L., Gomez-Alvarez, V., Revez, J., Palander, M., Antikainen, J., Kauppinen, A., Räsänen, P., Siponen, S., Nyholm, O., Kyyhkynen, A., Hakkarainen, S., Merentie, J., Pärnänen, M., Loginov, R., Ryu, H., Kuusi, M., Siitonen, A., Miettinen, I., Santo Domingo, J.W., Hänninen, M.L., Pitkänen, T. | Novel Microbiological and Spatial Statistical Methods to Improve Strength of Epidemiological Evidence in a Community-Wide Waterborne Outbreak | 2014 | PLoS One | [10.1371/journal.pone.0104713](https://doi.org/10.1371/journal.pone.0104713) | PRJNA235912 |
| 21 | Potgieter, S. C. & Pinto, A. J. | Reproducible microbial community dynamics of two drinking water systems treating similar source waters | 2019 | bioRxiv | [10.1021/acsestwater.1c00093](https://pubs.acs.org/doi/pdf/10.1021/acsestwater.1c00093) | PRJNA529765 |
| 22 | Liu, J., Zhao, R., Zhang, J., Zhang, G., Yu, K., Li, X., Li, B. | Occurrence and fate of ultramicrobacteria in a full-scale drinking water treatment plant | 2018 | Frontiers in Microbiology | [10.3389/fmicb.2018.02922](https://doi.org/10.3389/fmicb.2018.02922) | PRJNA481143 |
| 23 | Pereira R.P.A., Peplies J., Höfle M.G., Brettar I. | Bacterial community dynamics in a cooling tower with emphasis on pathogenic bacteria and Legionella species using universal and genus-specific deep sequencing | 2017 | Water Research | [10.1016/j.watres.2017.06.011](https://doi.org/10.1016/j.watres.2017.06.011) | PRJEB14855 |
| 24 | De Vera G.A., Gerrity D., Stoker M., Frehner W., Wert E.C. | Impact of upstream chlorination on filter performance and microbial community structure of GAC and anthracite biofilters | 2018 | Environmental Science: Water Research and Technology | [10.1039/c8ew00115d](https://doi.org/10.1039/c8ew00115d) | PRJNA450157 |
| 25 | de Vera G.A., Wert E.C. | Using discrete and online ATP measurements to evaluate regrowth potential following ozonation and (non)biological drinking water treatment | 2019 | Water Research | [10.1016/j.watres.2019.02.006](https://doi.org/10.1016/j.watres.2019.02.006) | PRJNA483220 |
| 26 | Gerrity D., Arnold M., Dickenson E., Moser D., Sackett J.D., Wert E.C. | Microbial community characterization of ozone-biofiltration systems in drinking water and potable reuse applications | 2018 | Water Research | [10.1016/j.watres.2018.02.023](https://doi.org/10.1016/j.watres.2018.02.023) | PRJEB22574 |
| 27 | Ahmed, W., Staley, C., Sadowsky, M. J., Gyawali, P., Sidhu, J. P. S., Palmer, A., Beale, D. J., Toze, S. | Toolbox Approaches Using Molecular Markers and 16S rRNA Gene Amplicon Data Sets for Identification of Fecal Pollution in Surface Water | 2015 | Applied and Environmental Microbiology | [10.1128/AEM.02032-15](https://doi.org/10.1128/AEM.02032-15) | PRJNA257794 |
