## Supplementary Information 2 for "Microbiomes in drinking water treatment and distribution: a meta-analysis from source to tap"

Table 2: Sequence Processing summary of all studies included in this analysis. Including the sequencing platform (HiSeq or MiSeq); Hypervariable region sequenced, and the type of data available. 3 taxonomic assignment methods were employed including Naïve Bayesian Classifier, Bayesian Least Common Ancestor (BLCA), both with the SILVA138 database. The 3^rd^ approach uses TaxAss- a freshwater database to assign taxonomy where SILVA138 fails.

| **ID** | **Studies** | **Illumina Sequencing Platform** | **Hyper-variable 16S region** | **Read Type** | **QIIME2 Processing Pipeline** | **Demultiplexed Reads** | **ASVs to sequence level**  **NBC + SILVA 138** | **ASVs to sequence level BLCA + SILVA138** | **ASVs to sequence level**  **TaxAss** |
| --- | --- | --- | --- | --- | --- | --- | --- | --- | --- |
| 1 | PRJNA399213 | HiSeq | V3-V4 | Paired-end | DEBLUR | 2.23E+06 | 802485 | 801824 | 741388 |
| 2 | PRJNA451227 | HiSeq | V3-V4 | Merged-Paired-end | CUTADAPT/DEBLUR | 8.62E+05 | 505602 | 613471 | 613587 |
| 3 4 | PRJNA393048 PRJNA397332 | HiSeq | V4-V5 | Paired-end | DEBLUR | 6.79E+06 | 2354346 | 2311026 | 2348845 |
| 5 6 | PRJNA347544 PRJNA479666 | MiSeq | V1-V3 | Paired-end | DEBLUR | 3.67E+06 | 1578215 | 1541253 | 1577355 |
| 7 | PRJNA433427 | MiSeq | V3 | Paired-end | DADA2 | 1.27E+07 | 3589723 | 3586612 | 3562771 |
| 8 9 10 11 12 13 | PRJEB24989 PRJNA486727 PRJNA494637 PRJUOW1 PRJNA486366 PRJNA487710 | MiSeq | V3-V4 | Paired-end | DEBLUR | 6.22E+07 | 20319897 | 2718581 | 20278970 |
| 14 | QIITA 10251 | MiSeq | V4 | Merged-Paired-end | Import BIOM | 4.12E+07 | 22059127 | 21894682 | 22015786 |
| 15 16 17 18 19 20 21 | PRJEB7435 PRJNA287840 PRJDB5406 PRJNA283789 PRJNA445682 PRJNA235912 PRJNA529765 | MiSeq | V4 | Paired-end | DADA2 | 1.82E+08 | 63928949 | 62835733 | 62253019 |
| 22 | PRJNA481143 | HiSeq | V4 | Merged-Paired-end | CUTADAPT/DADA2 | 2.32E+06 | 2241610 | 2221244 | 2226277 |
| 23 24 25 | PRJEB14855 PRJNA450157 PRJNA483220 | MiSeq | V4-V5 | Paired-end | DEBLUR | 2.05E+06 | 1245816 | 1240790 | 1242664 |
| 26 | PRJEB22574 | MiSeq | V4-V5 | Merged-Paired-end | CUTADAPT/DEBLUR | 1.32E+06 | 666352 | 944538 | 944117 |
| 27 | PRJNA257794 | MiSeq | V5-V6 | Paired-end | DEBLUR | 1.52E+07 | 7626045 | 7613247 | 7603371 |
| **TOTAL** |  |  |  |  |  | **3.3250E+08** | **2.0199E+07** | **1.0832E+08** | **1.2541E+08** |
| **Unique Taxa** |  |  |  |  |  |  |  | **22574** | **4858** |
