## Supplementary Information 3 for "Microbiomes in drinking water treatment and distribution: a meta-analysis from source to tap"

*The 25 most abundant genera for all abundance tables (grouped by their sequencing instrument and the hypervariable region sequenced) annotated: a) prior to collation and extraction of full-length sequences from the SILVA database and b) after collation. Results of Mantel and Procrustes tests of bray distances between individual and collated tables are shown below plots.*

### 1. MiSeq V1-V3

a)

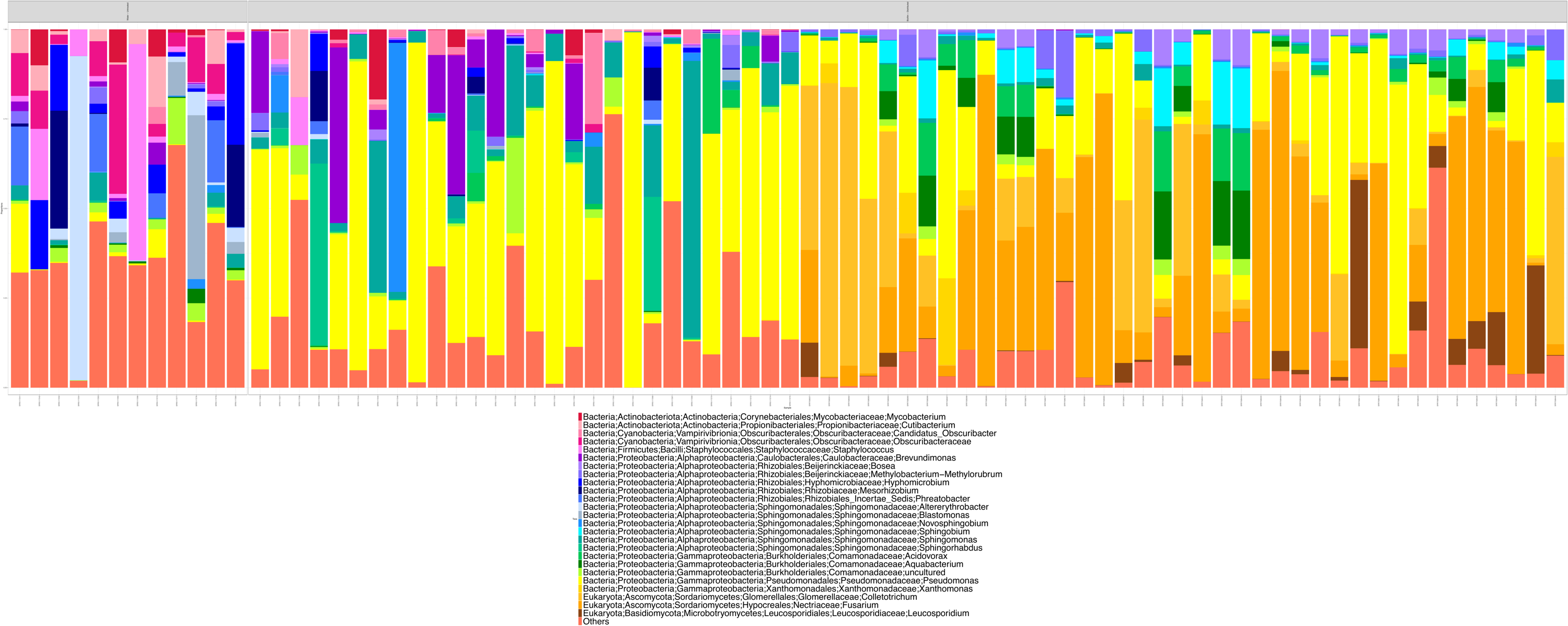

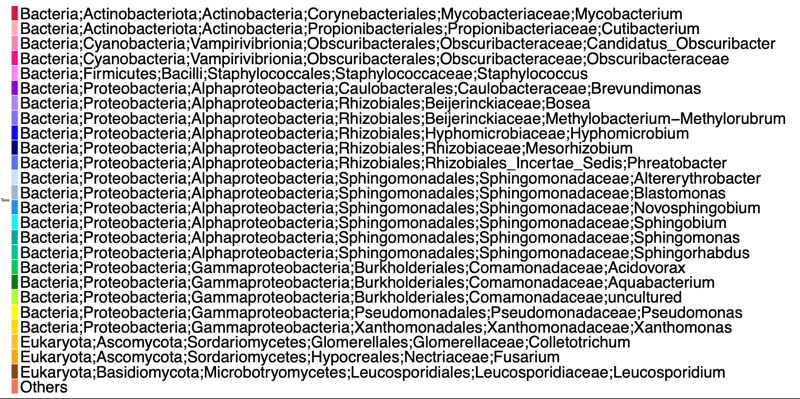

b)

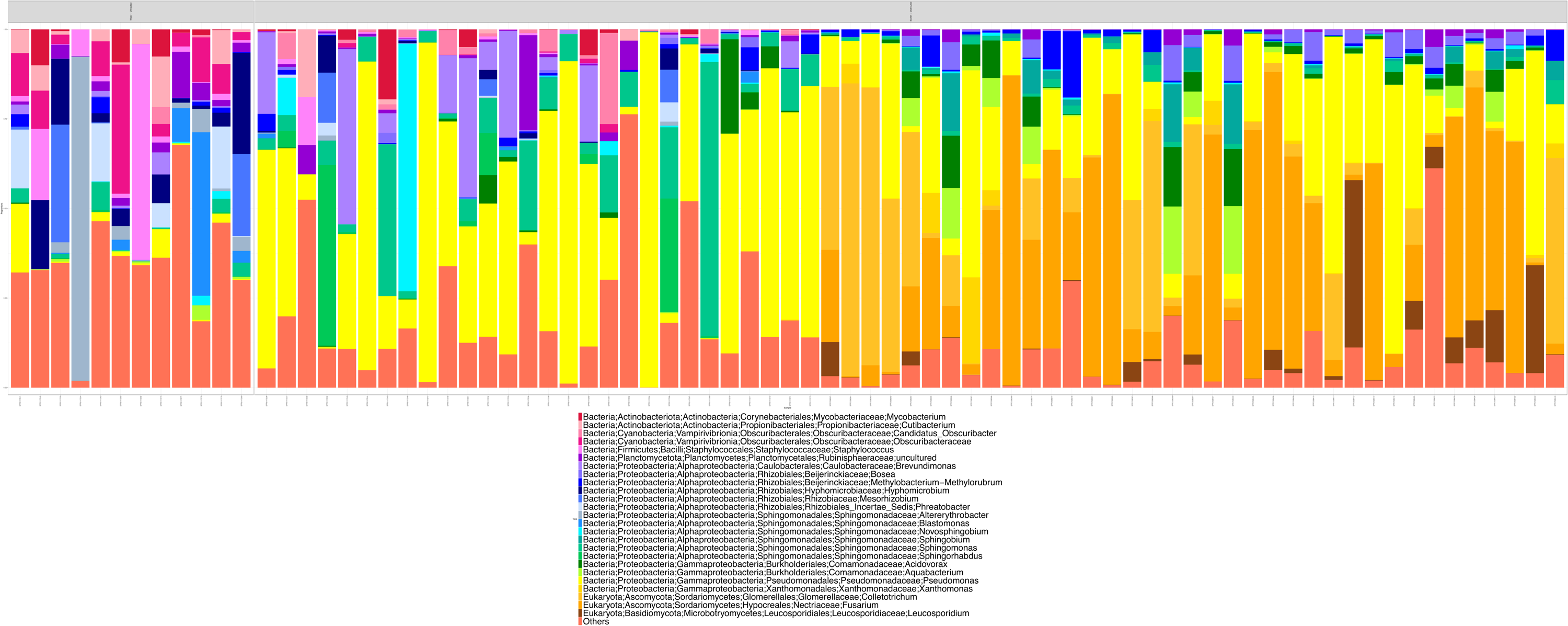

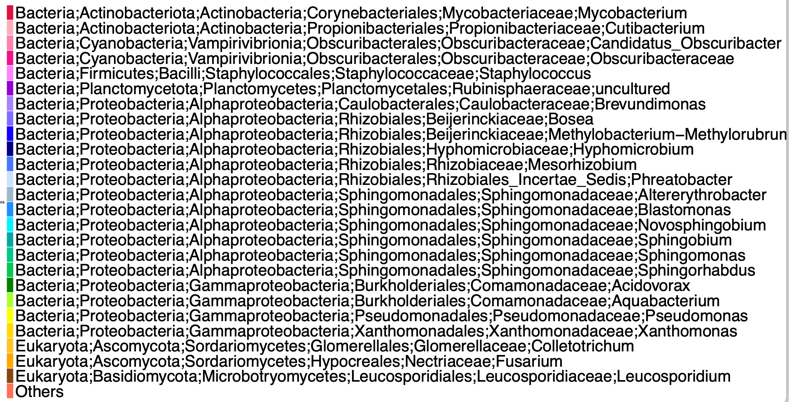

r = 0.9030119 | p = 0.0001 | Sum of Squares **=** 0.5978 | Correlation = 0.6342 | p = 0.001

### 2. MiSeq V3

a)

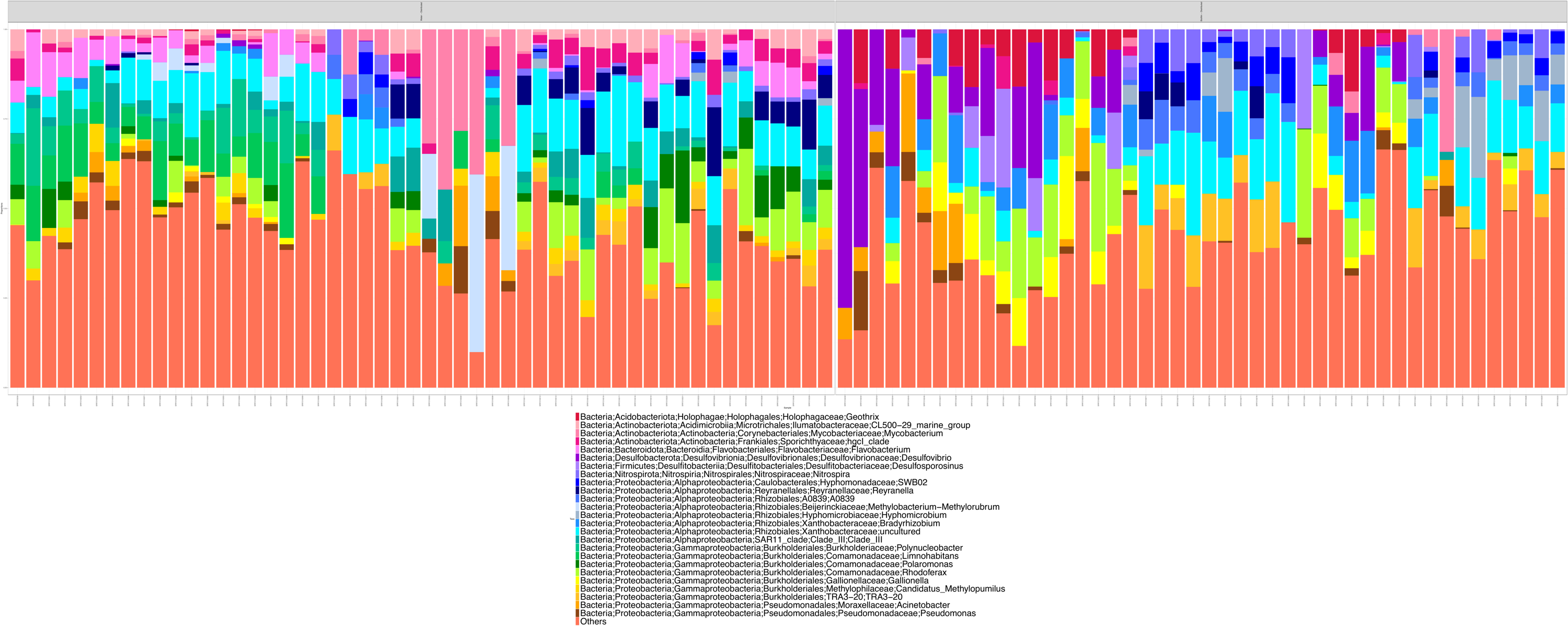

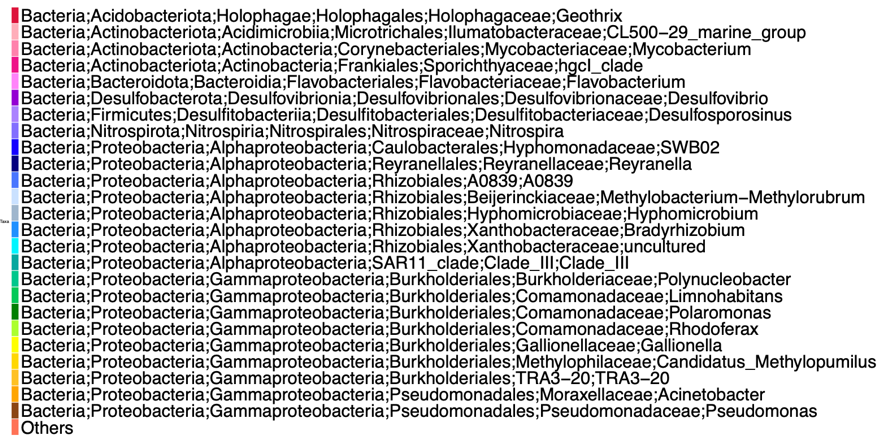

b)

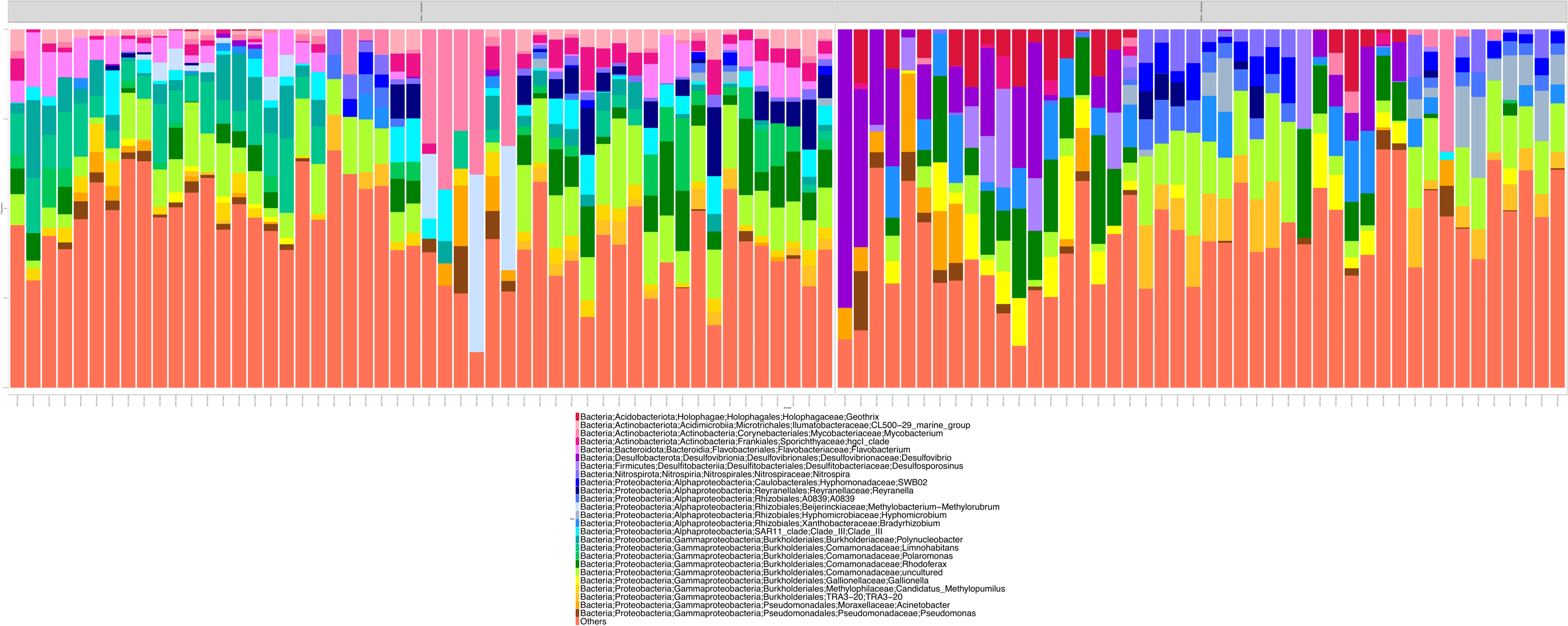

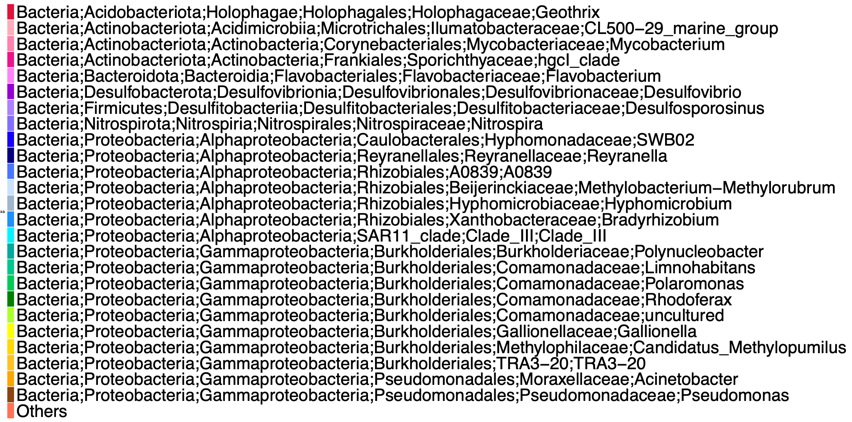

r = 0.041207 | p = 0.0012 | Sum of Squares= 0.97 | Correlation = 0.17 | p = 0.12

### 3. MiSeq V4 (separated into 2 biom files: i) and ii) due to large dataset)

i) a)

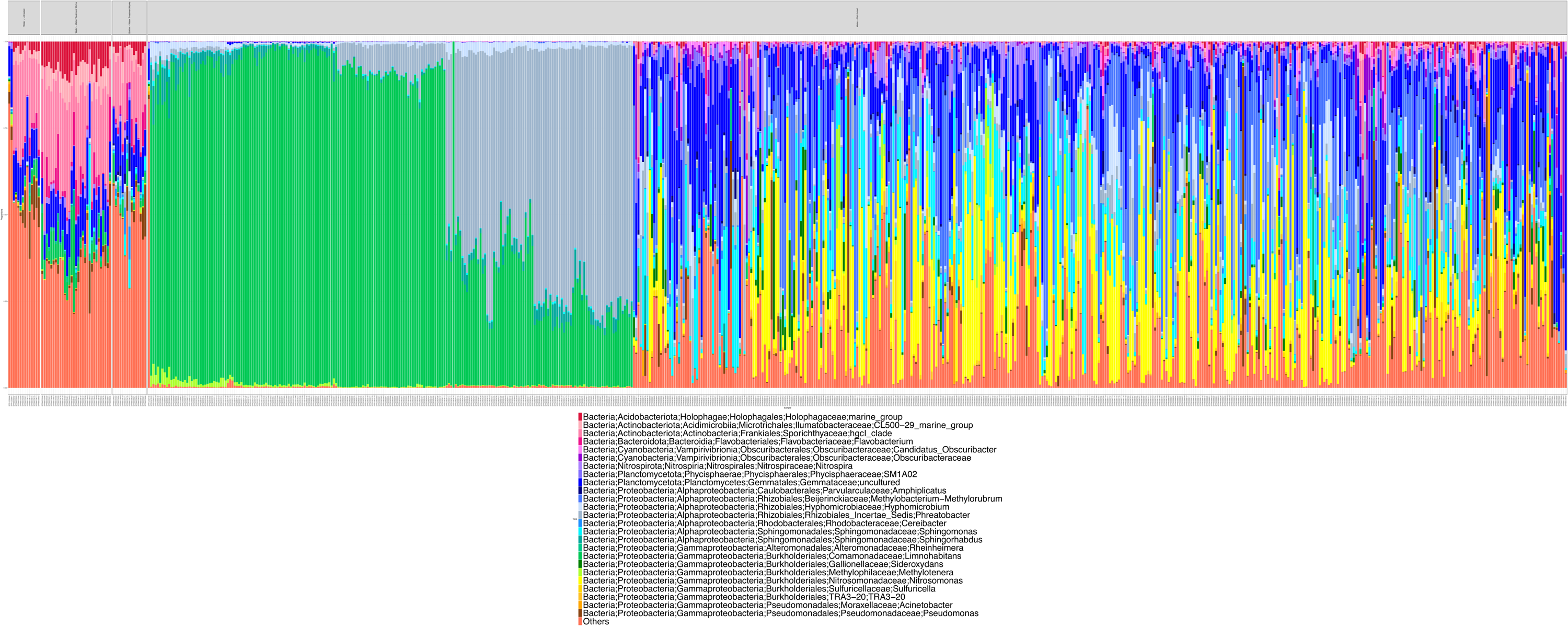

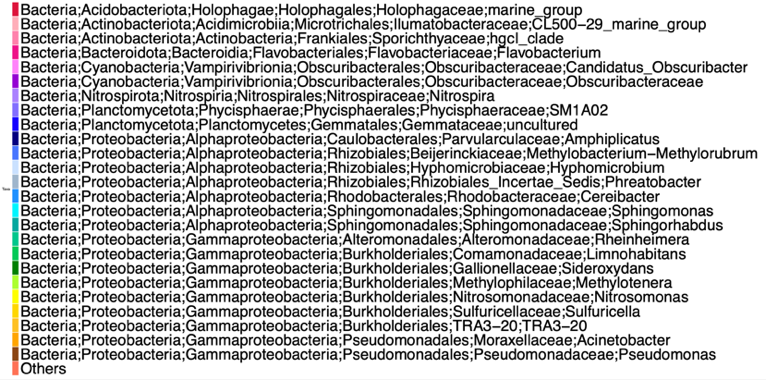

b)

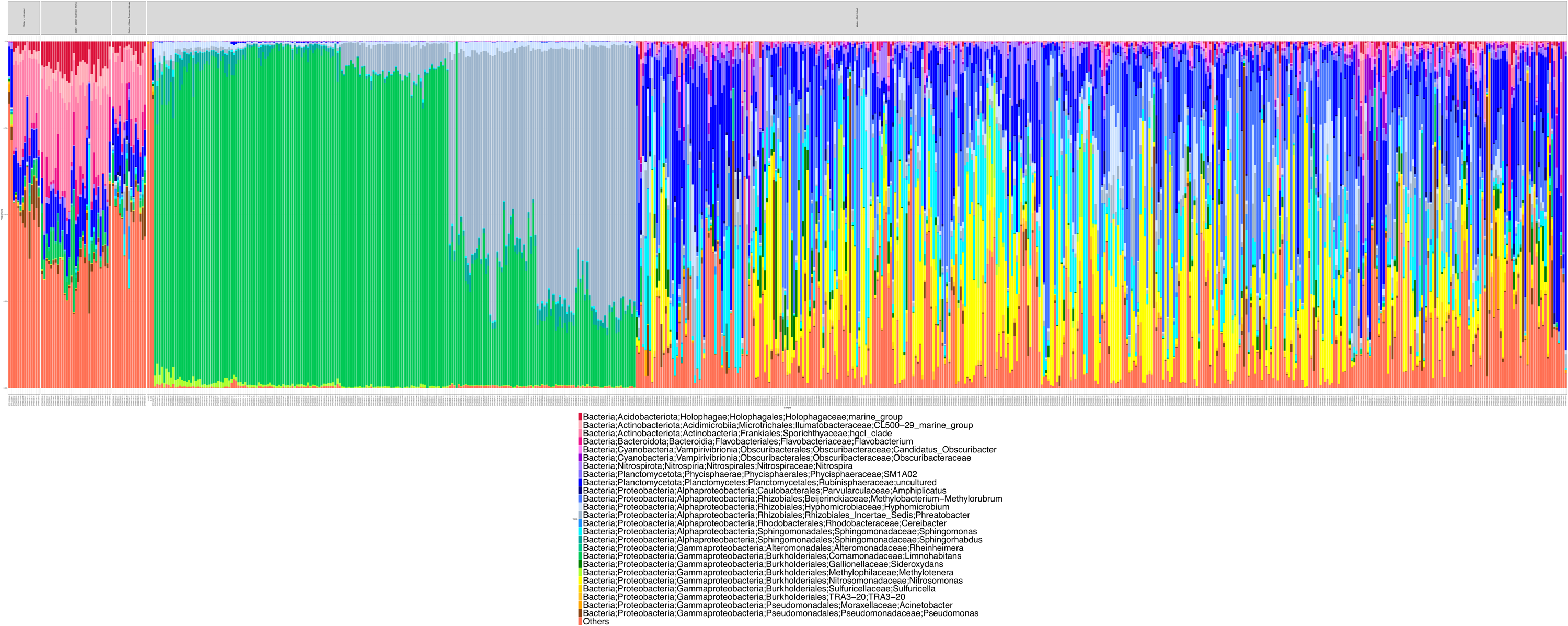

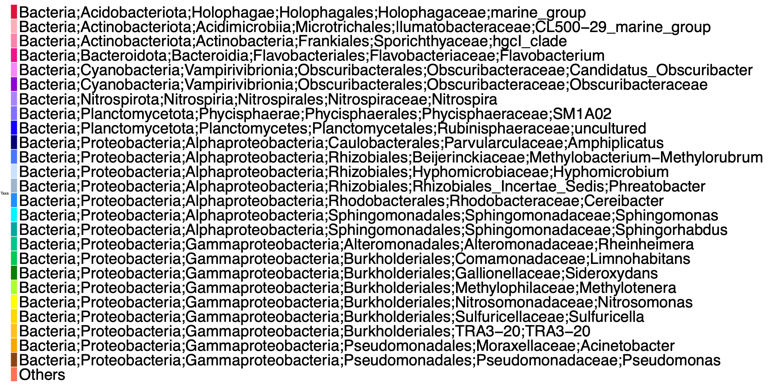

r = 0.9755339 | p = 0.0001 | Sum of Squares = 0.9842 | Correlation = 0.1255 | p = 0.001

ii) a)

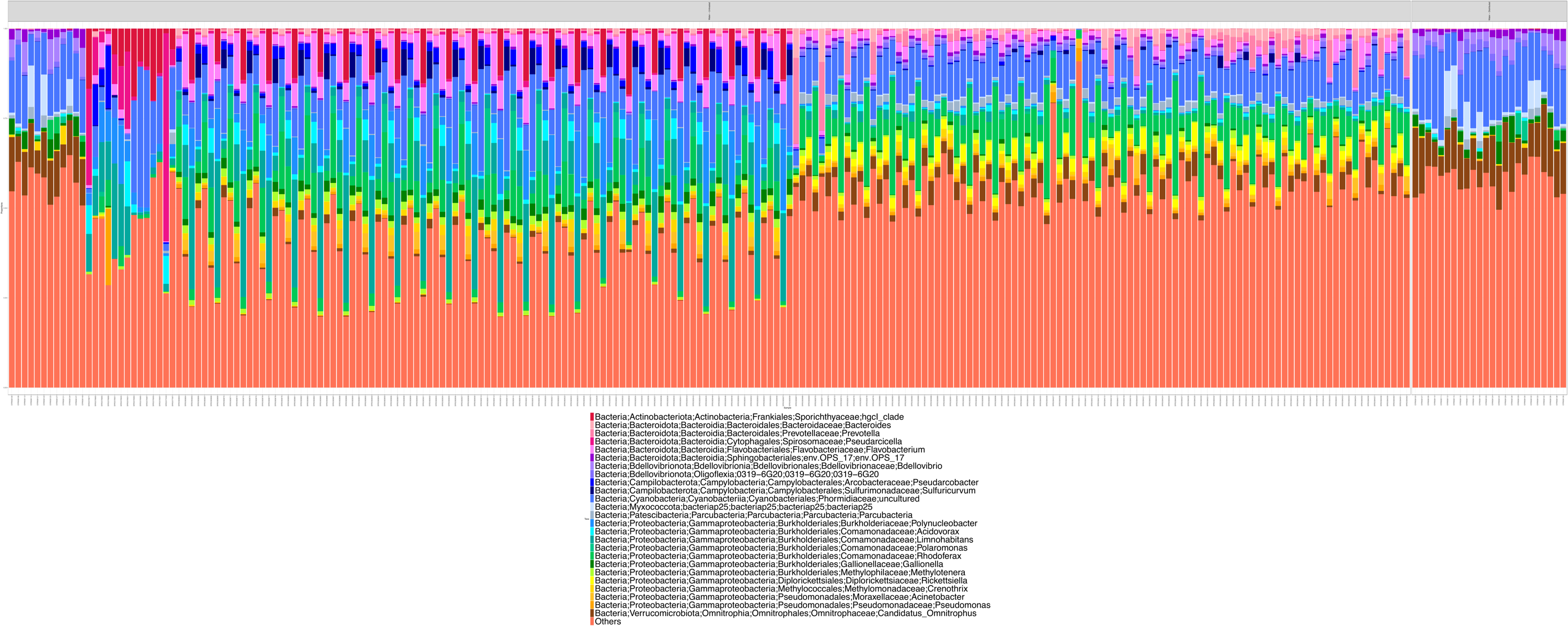

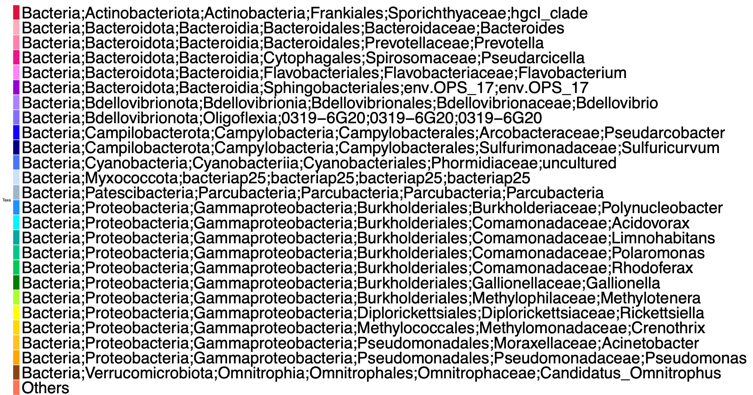

b)

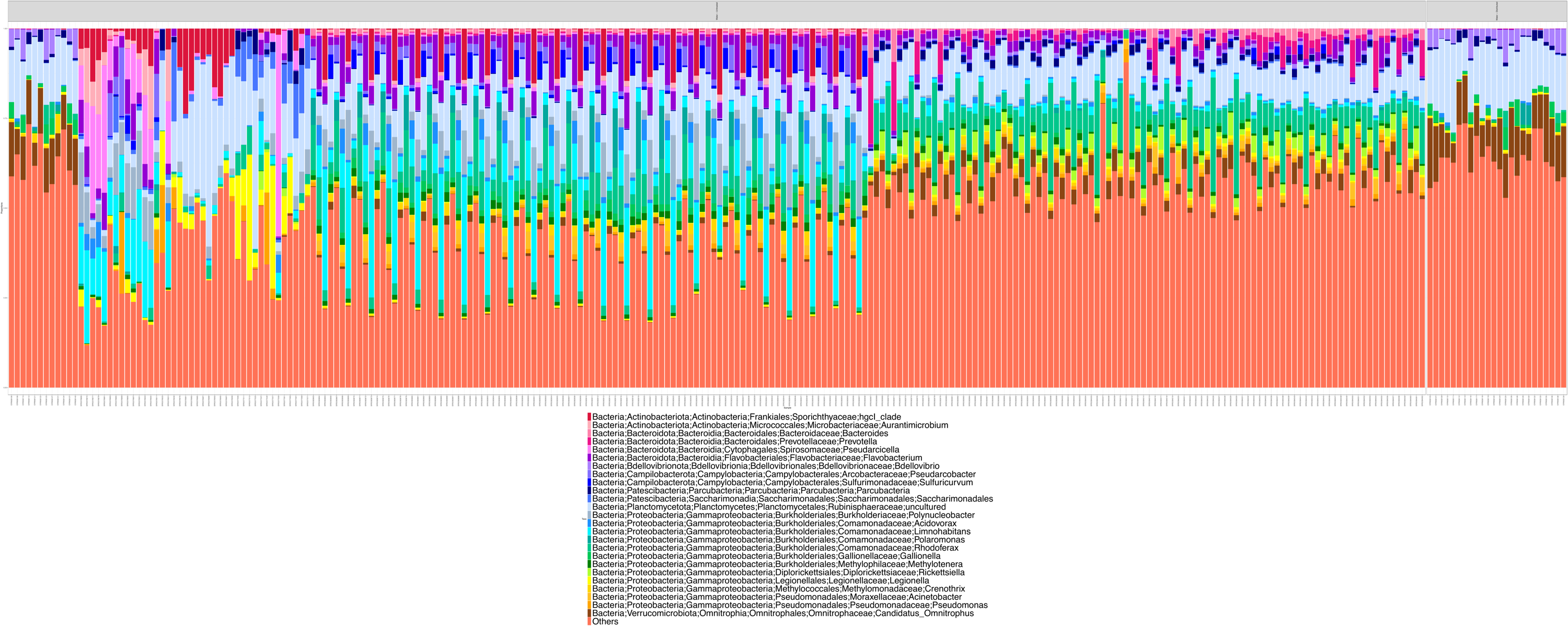

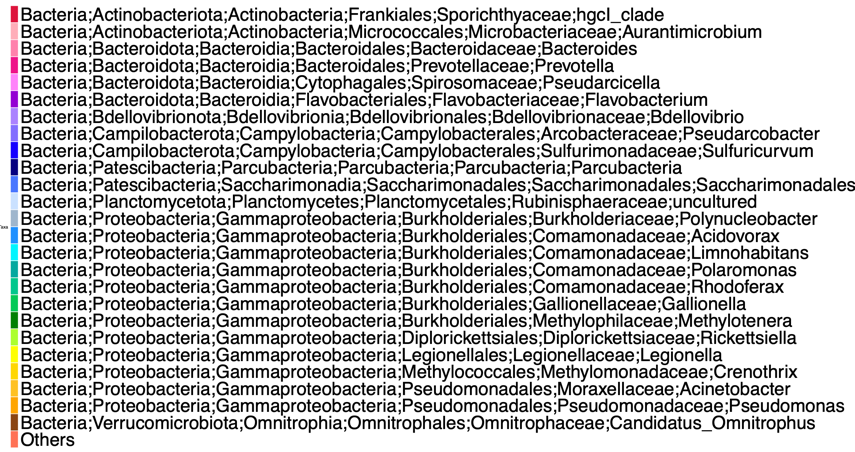

r = 0.9291108 | p = 0.0001 | Sum of Squares = 0.7131 |Correlation = 0.5356 | p = 0.001

### 4. MiSeq V4 (QIITA Data)

a)

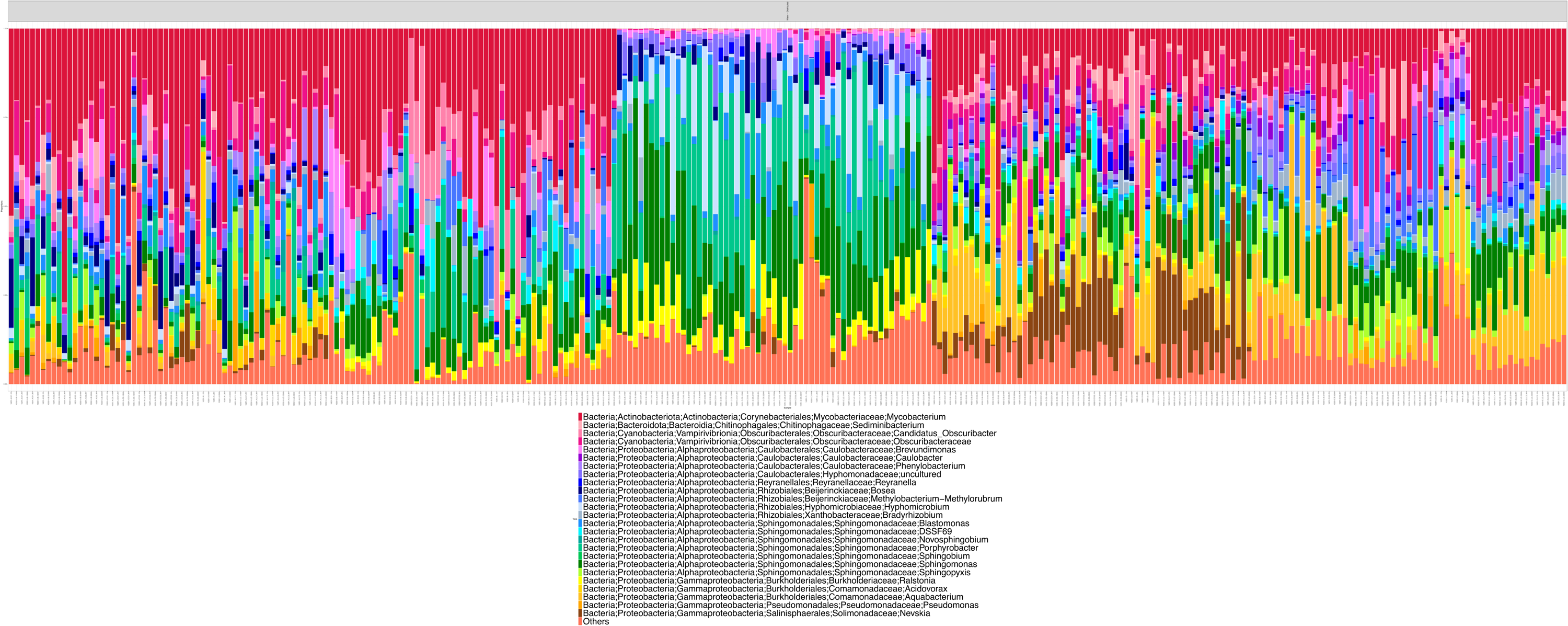

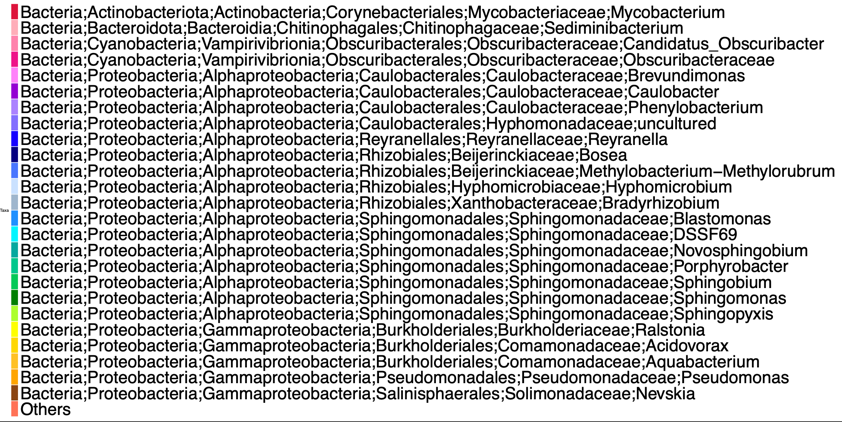

b)

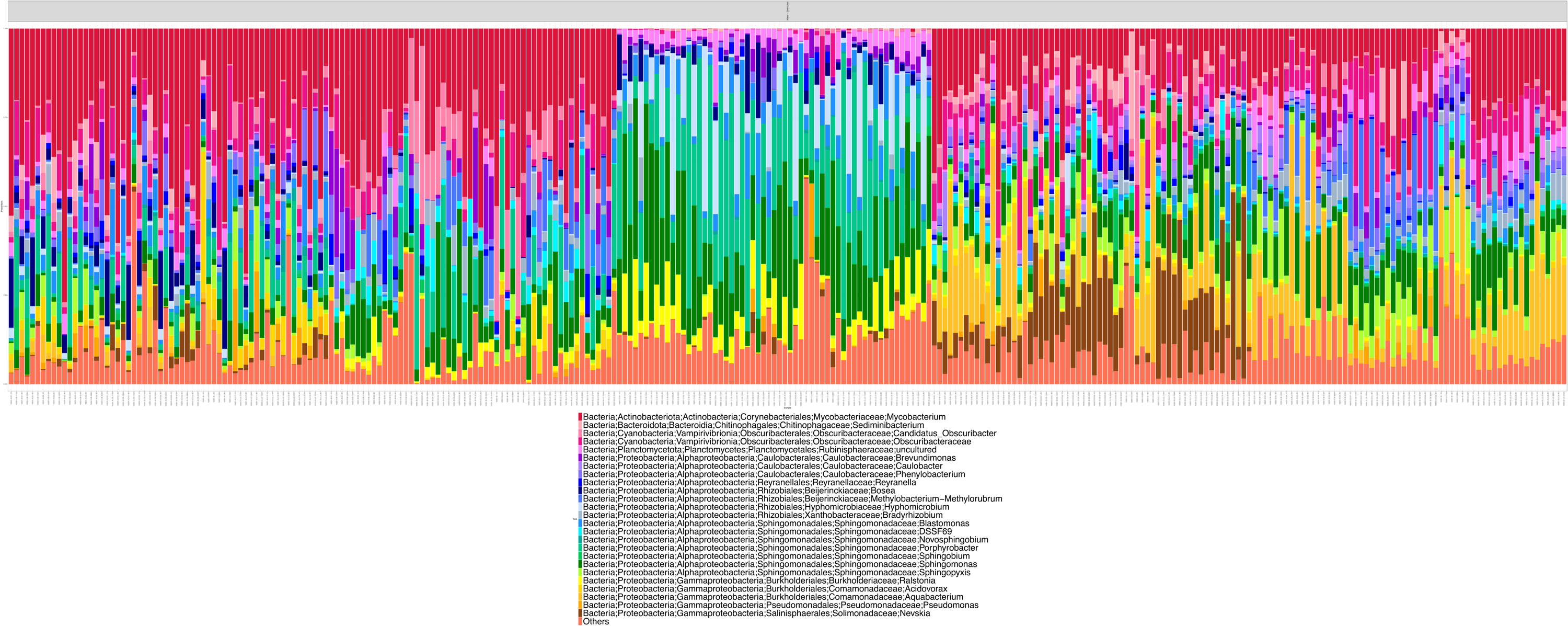

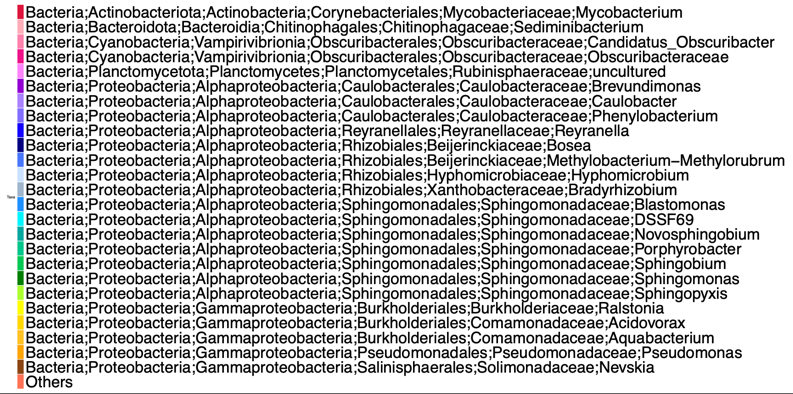

r = 0.935775 | p = 0.0001 | Sum of Squares = 0.2795 | Correlation = 0.8488 | p = 0.001

### 5. MiSeq V4-V5

a)

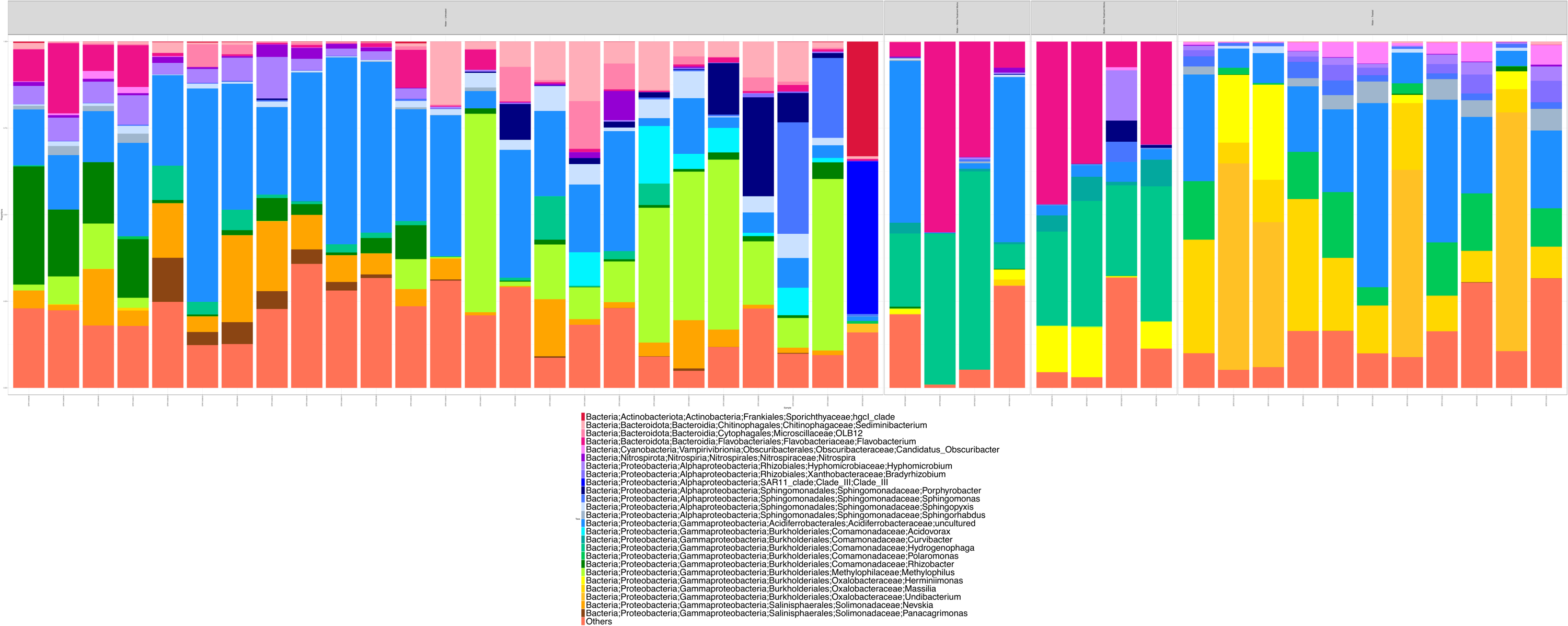

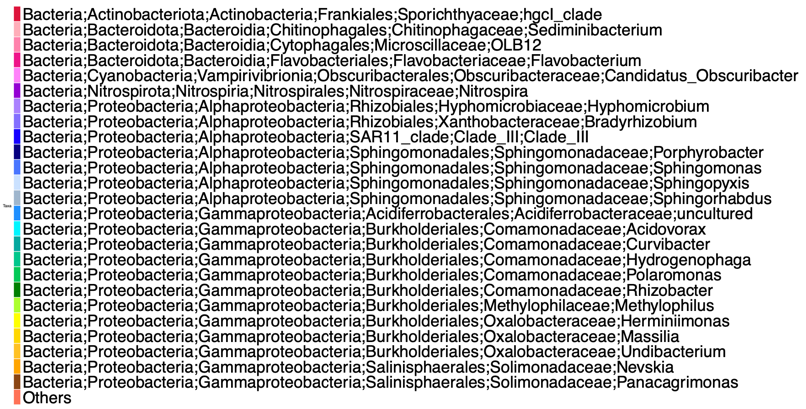

b)

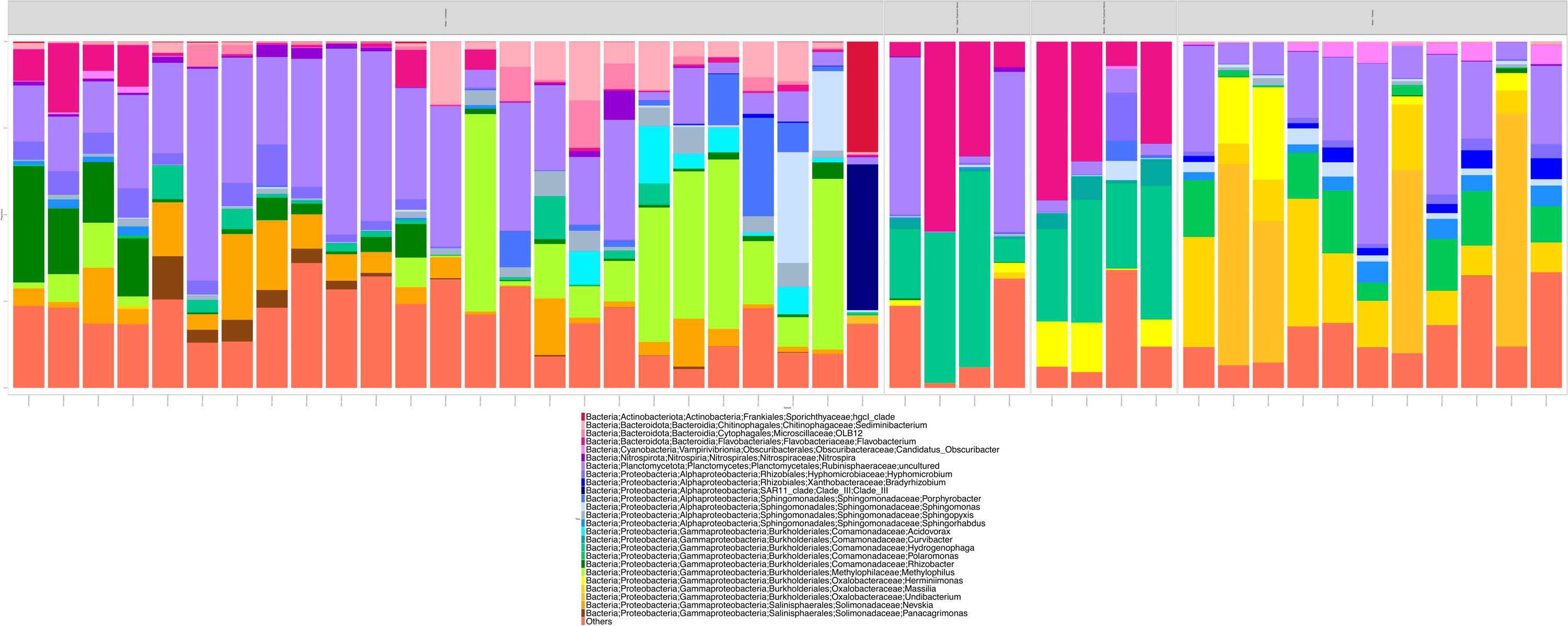

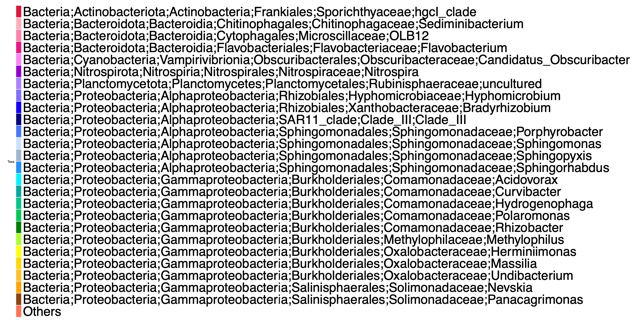

r = 0.9867119 | p = 0.0001 | Sum of Squares **=** 0.488 | Correlation = 0.7155 | p = 0.001

### 6. MiSeq V4-V5 Merged-Paired-End

a)

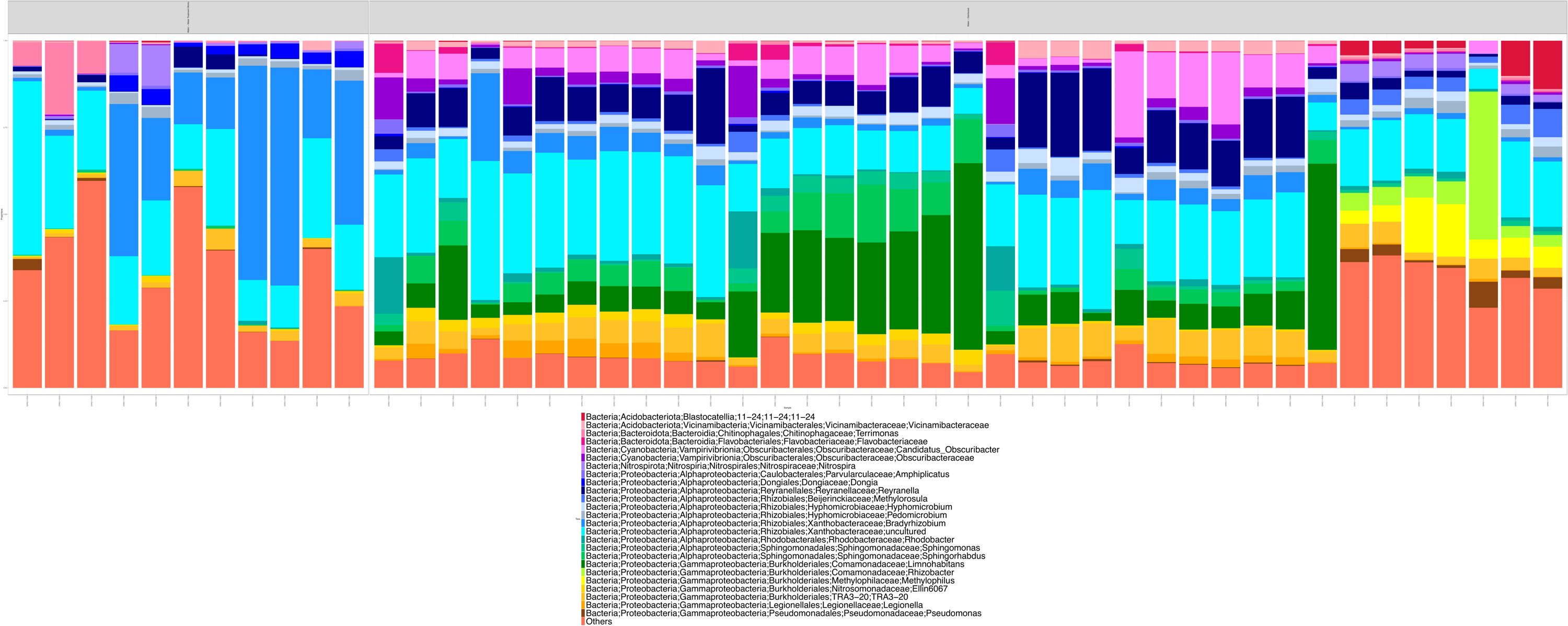

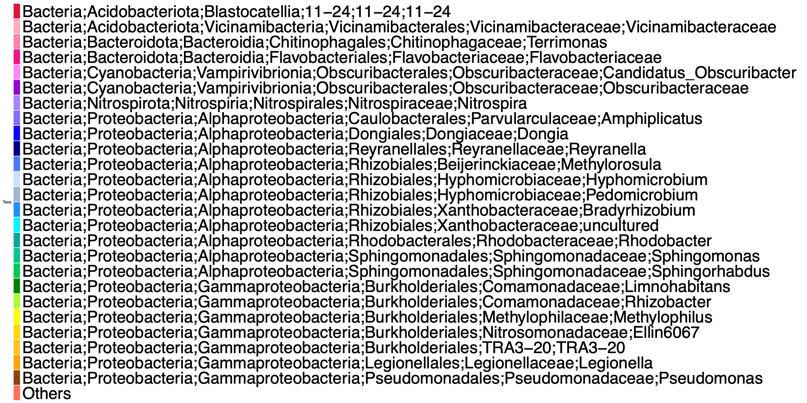
b)

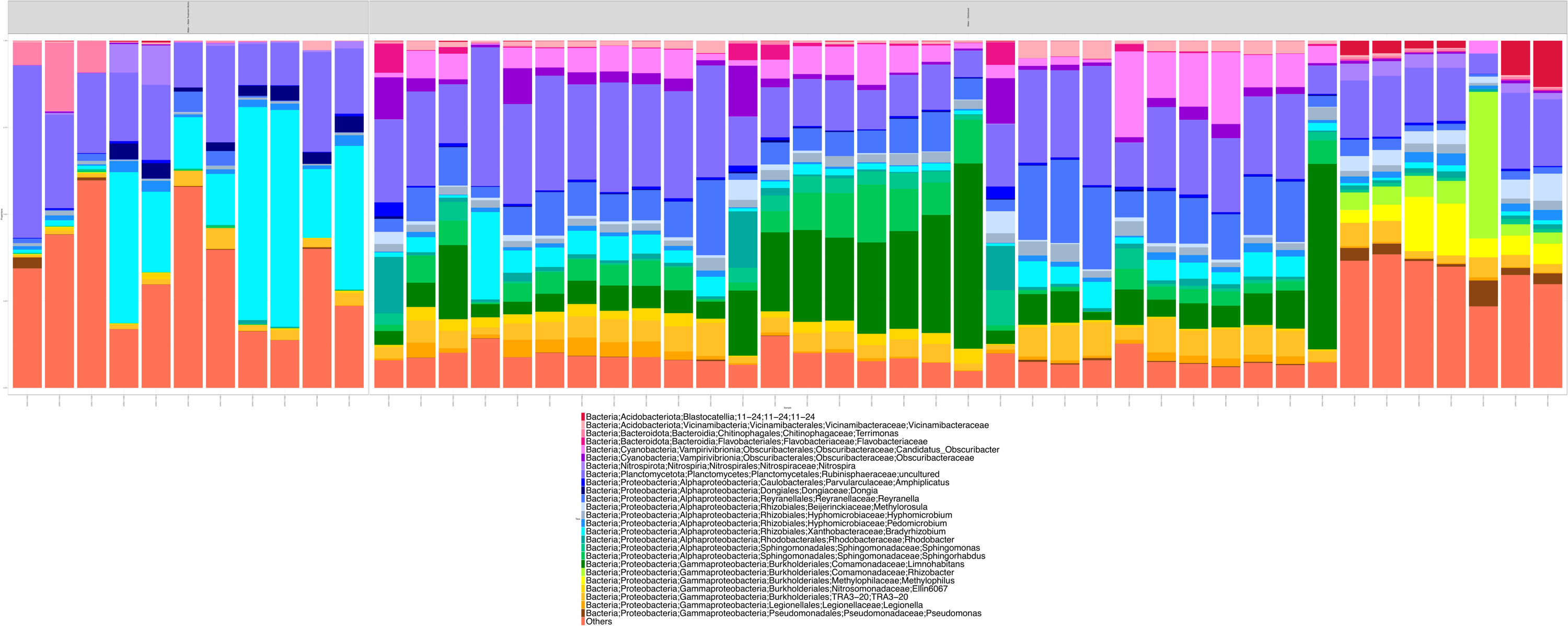

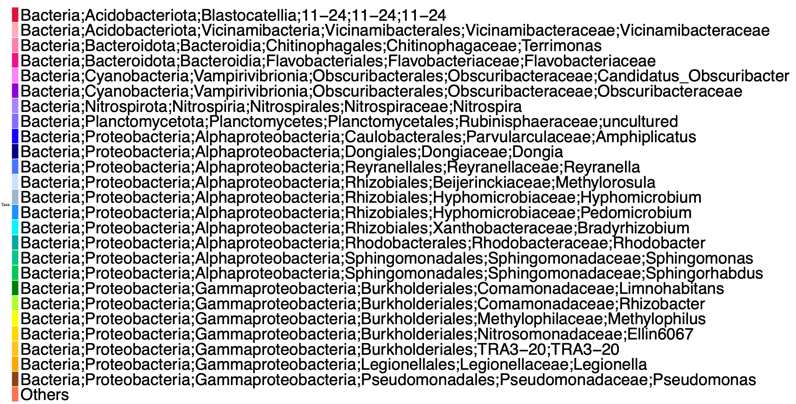

r = 0.9915712 | p = 0.0001 | Sum of Squares **=** 0.4914 |Correlation = 0.7131 | p = 0.001

### 7. MiSeq V5-V6

a)

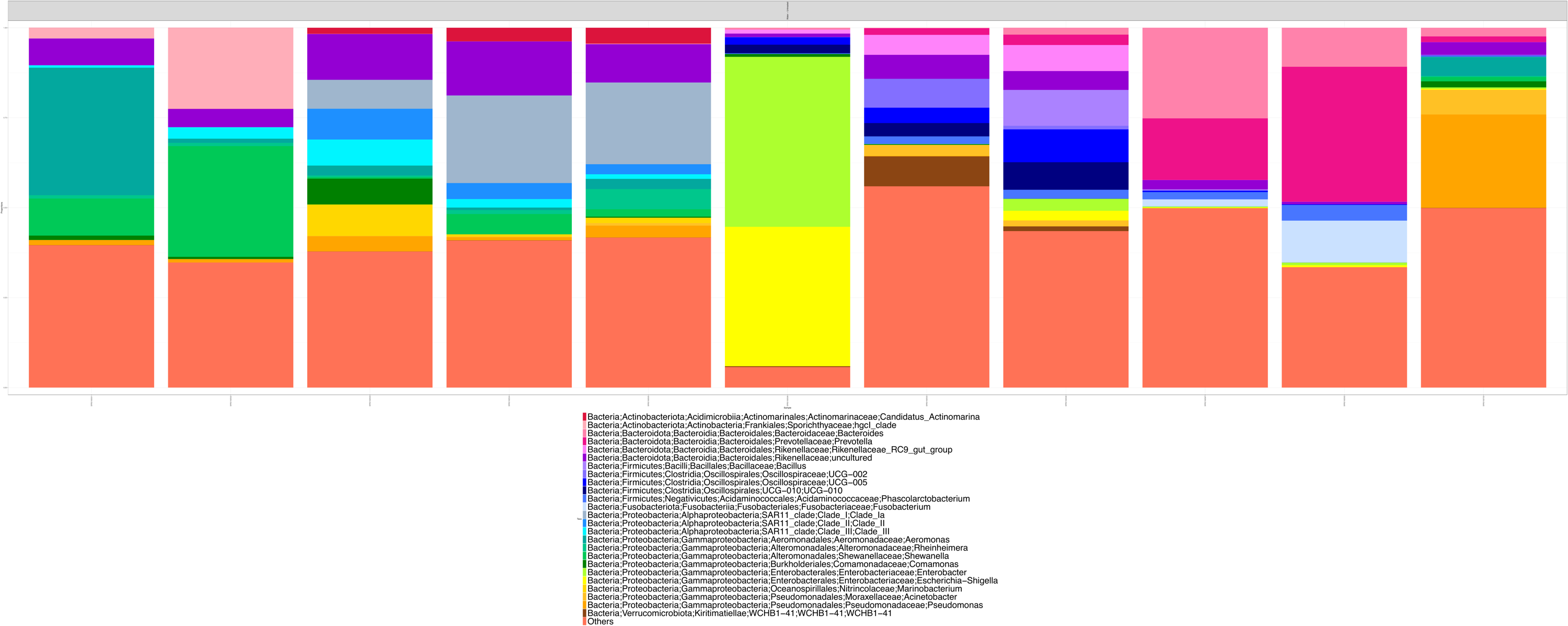

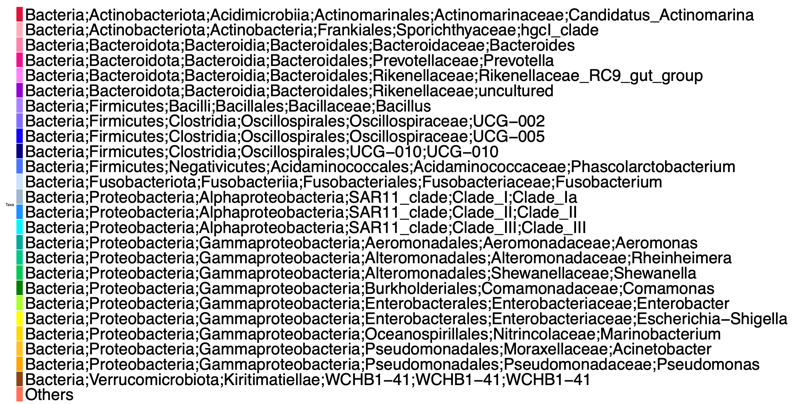

b)

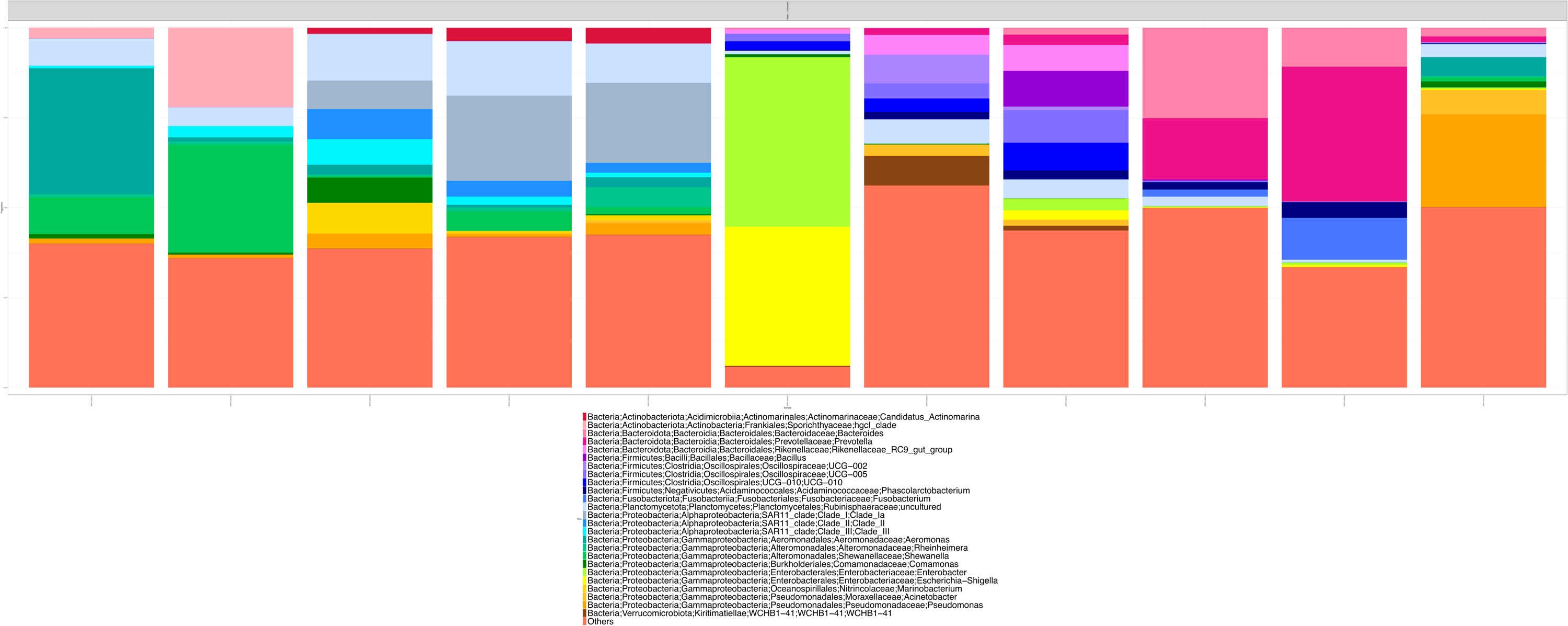

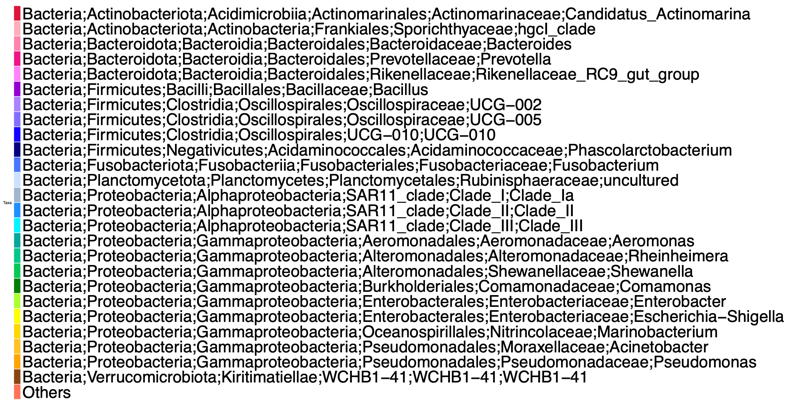

r = 0.9321193 | p = 0.0001 | Sum of Squares **=** 0.3037 | Correlation = 0.8345 | p = 0.001

### 8. MiSeq V3-V4

a)

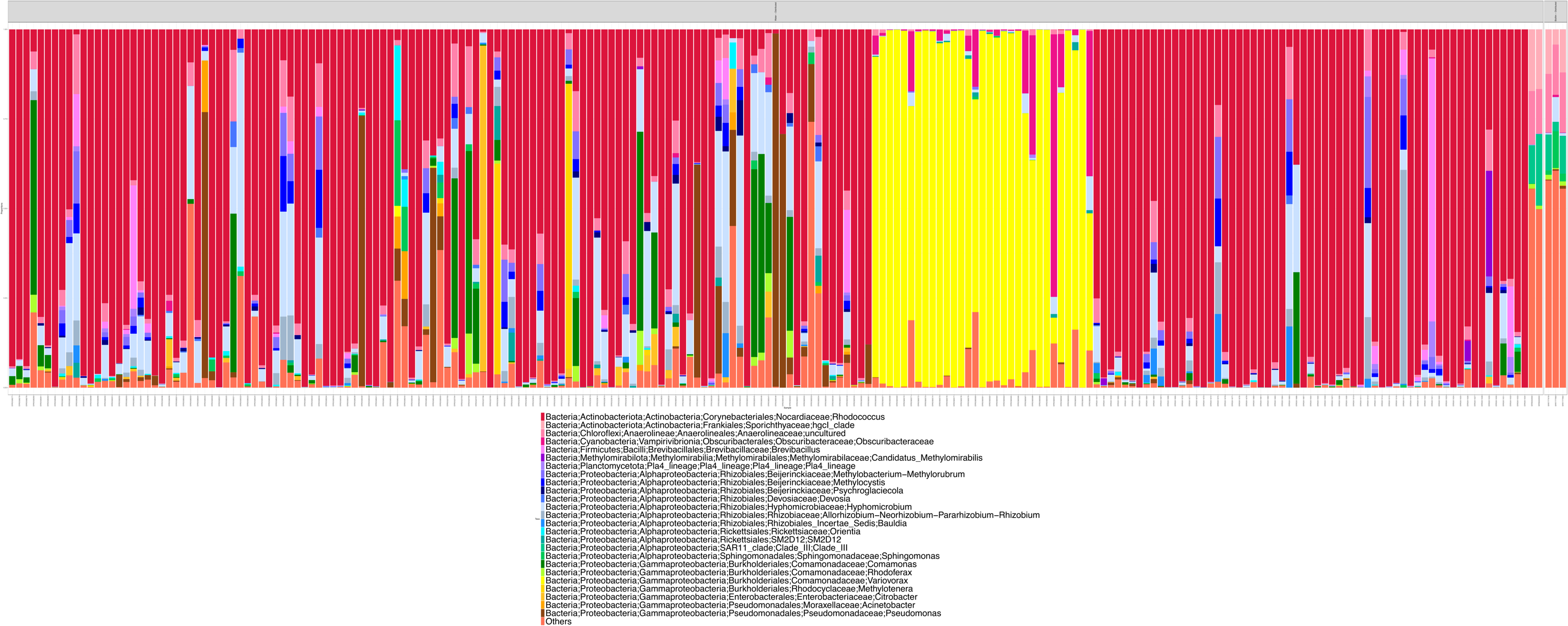

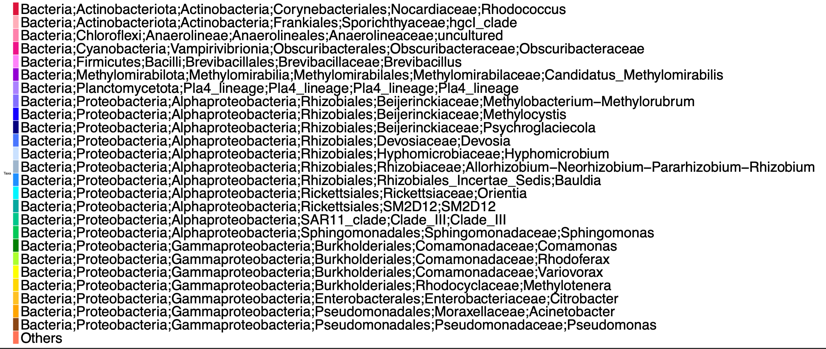

b)

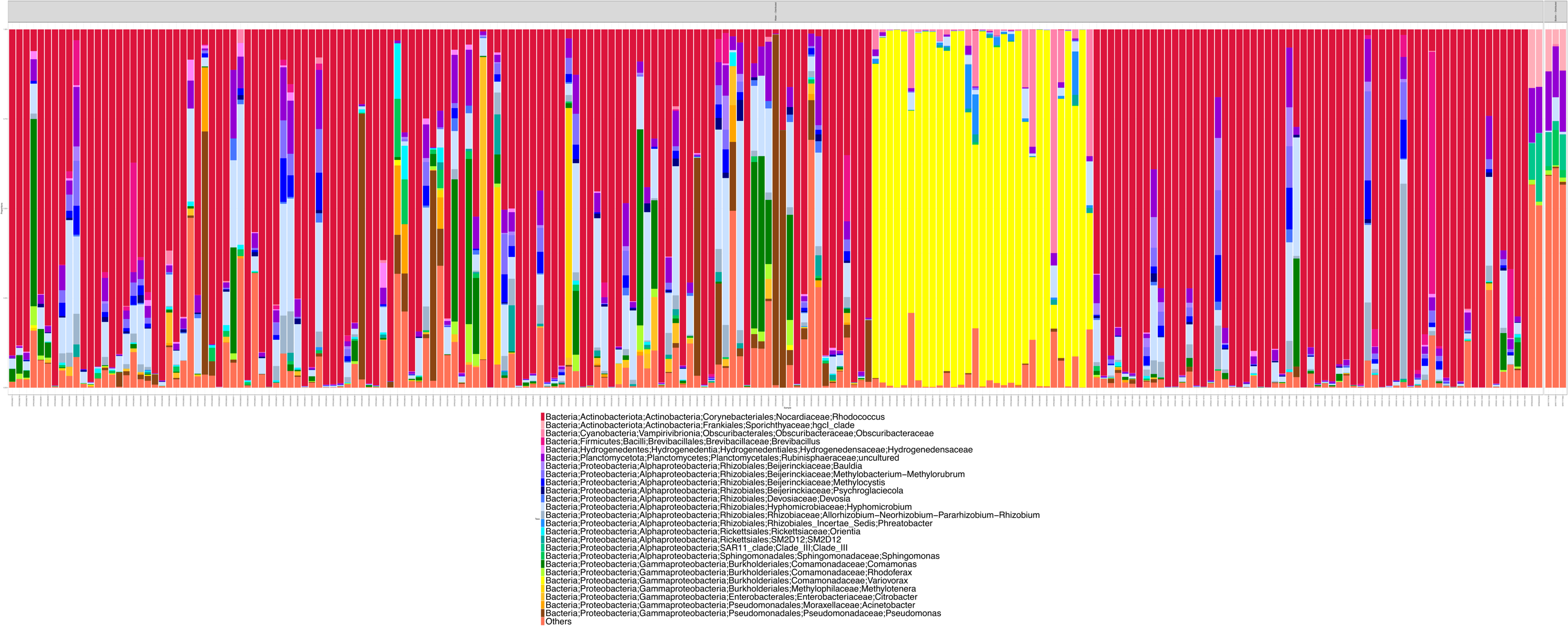

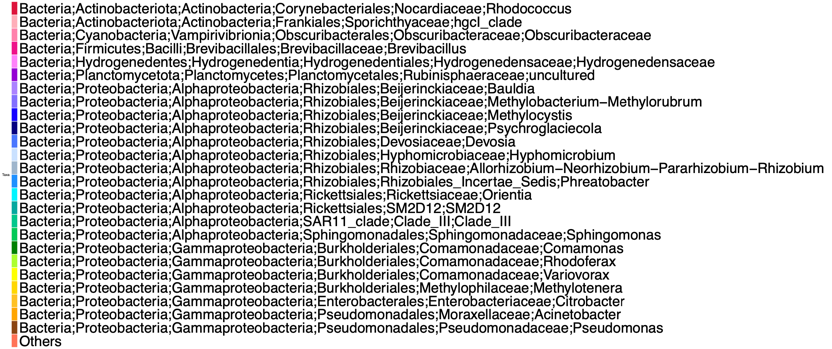

r = 0.5494883 | p = 0.0001 | Sum of Squares = 0.7283 | Correlation = 0.5213 | p = 0.001

### 9. HiSeq V3-V4

a)

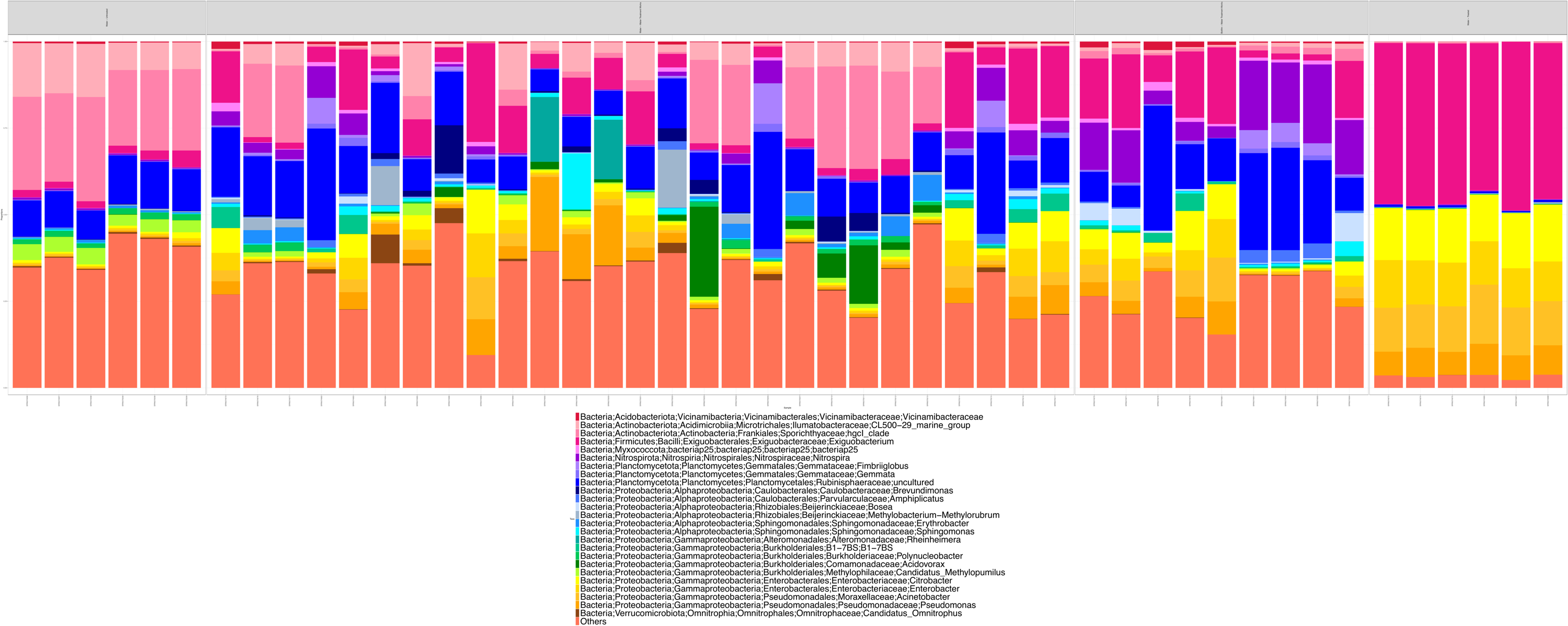

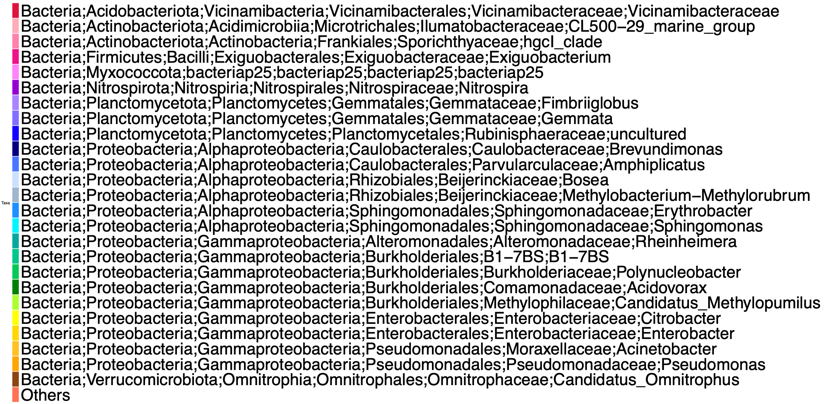

b)

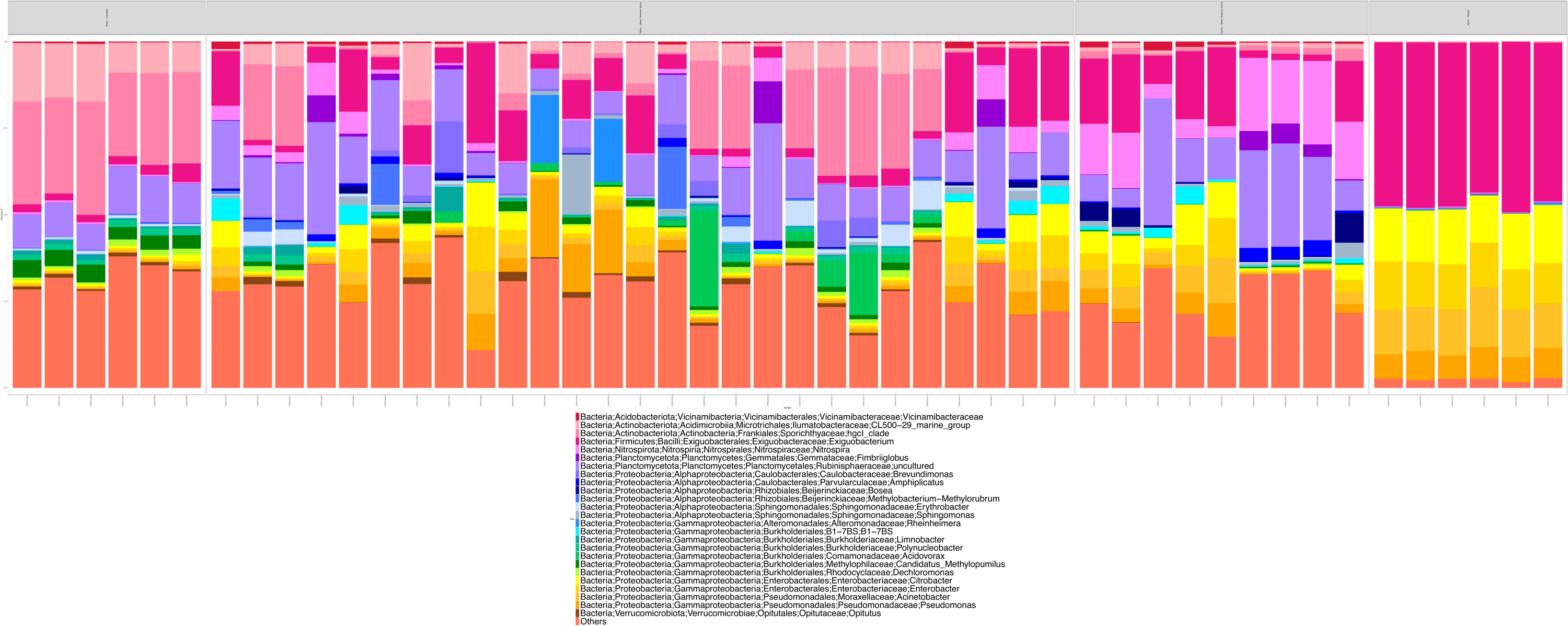

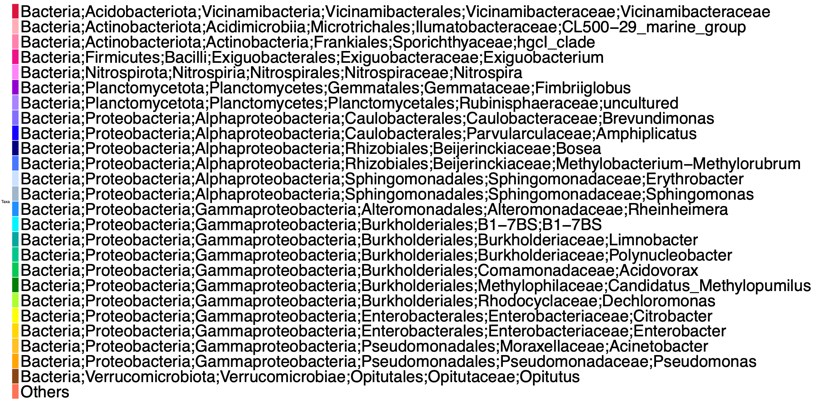
 r = 0.9685338 | p = 0.0001000 | Sum of Squares = 0.1529 | Correlation= 0.9204 | p = 0.001

### 10. HiSeq V3-V4 Merged-Paired-End

a)

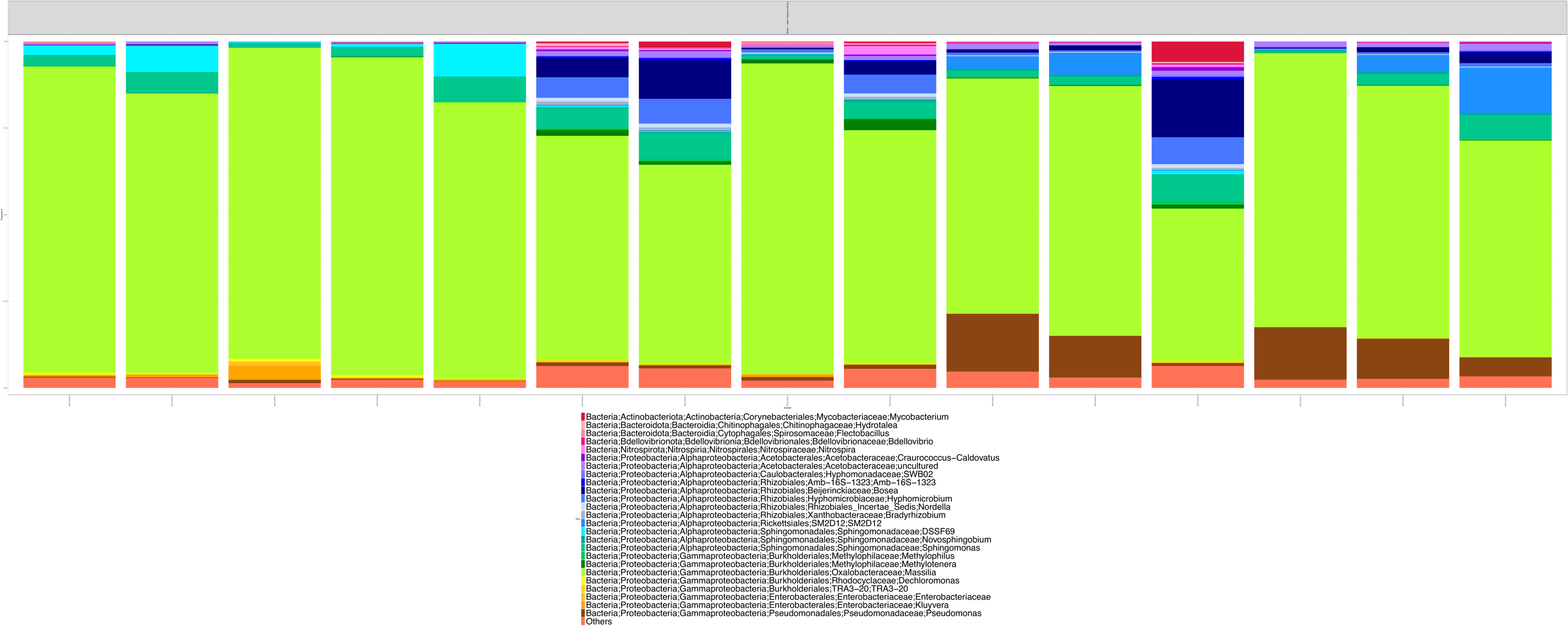

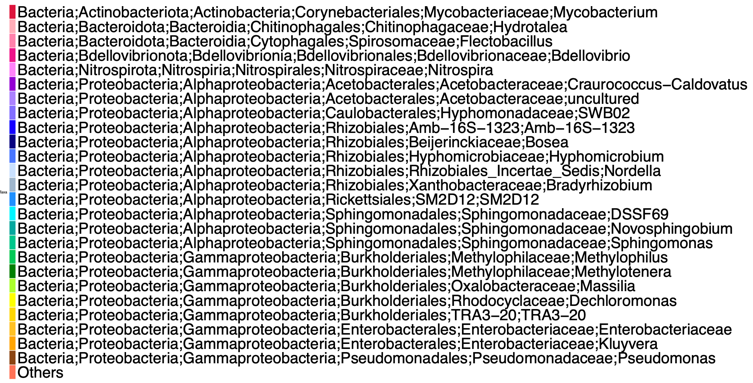

b)

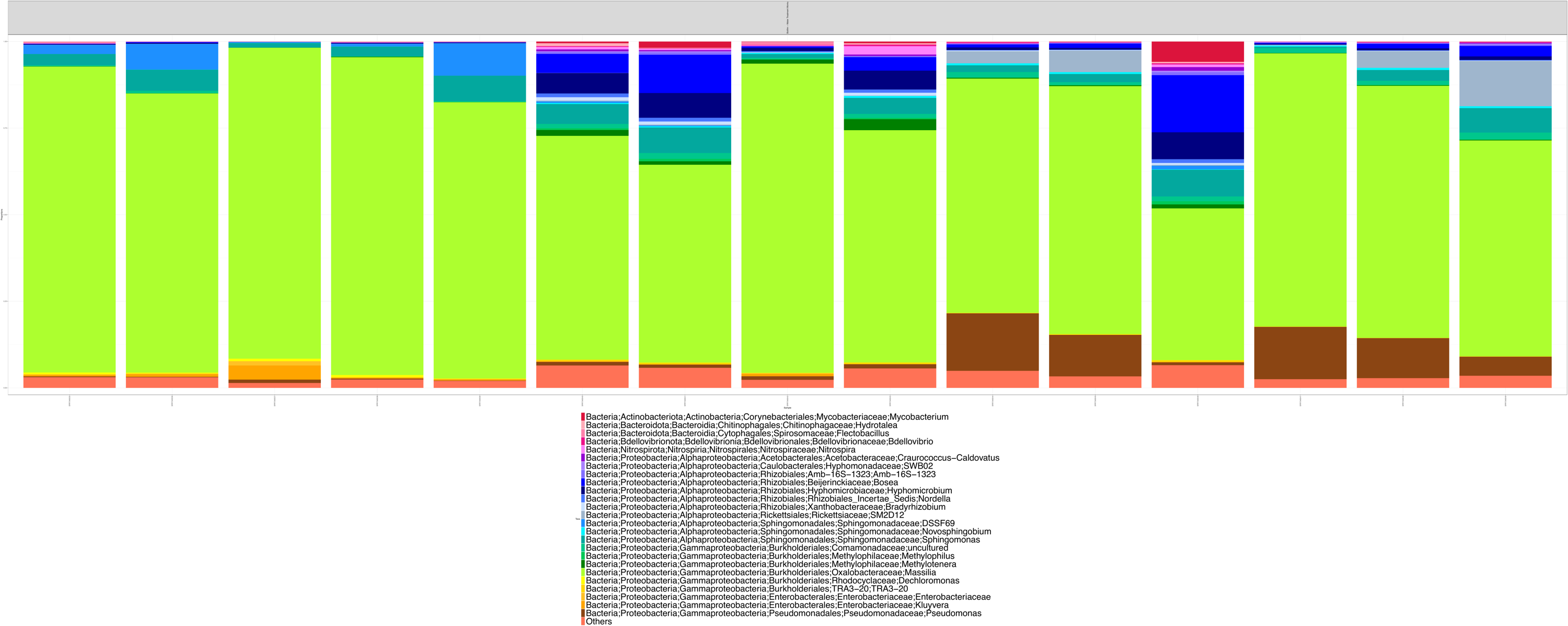

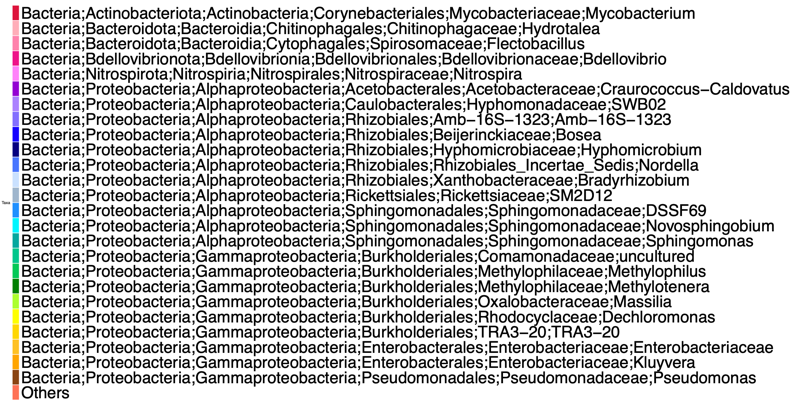

r = 0.9959586| p = 0.0001 | **Sum of Squares =** 0.1168 | Correlation = 0.9398 | p = 0.001

### 11. HiSeq V4

a)

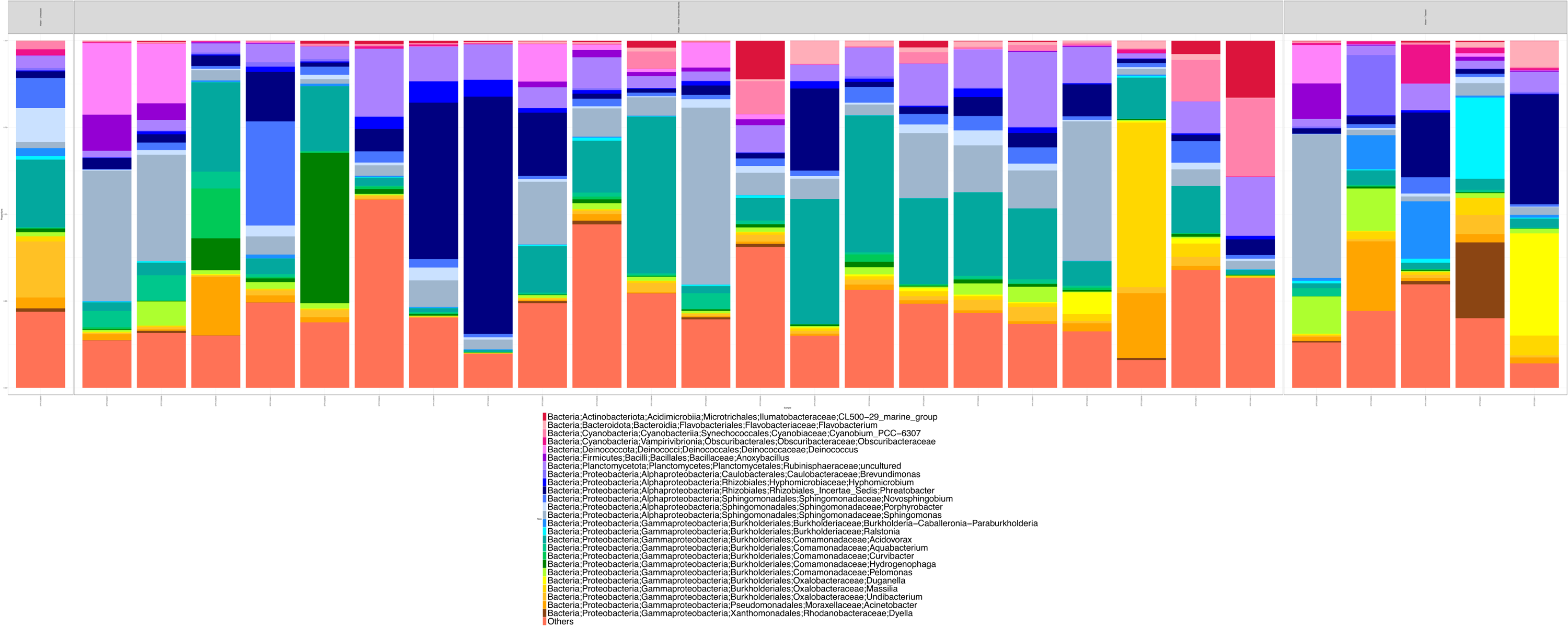

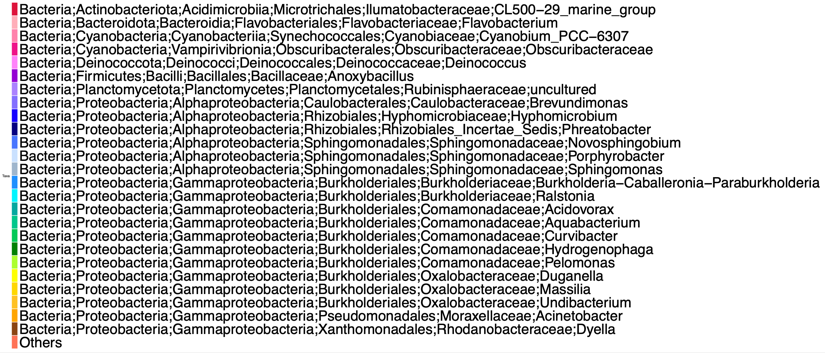

b)

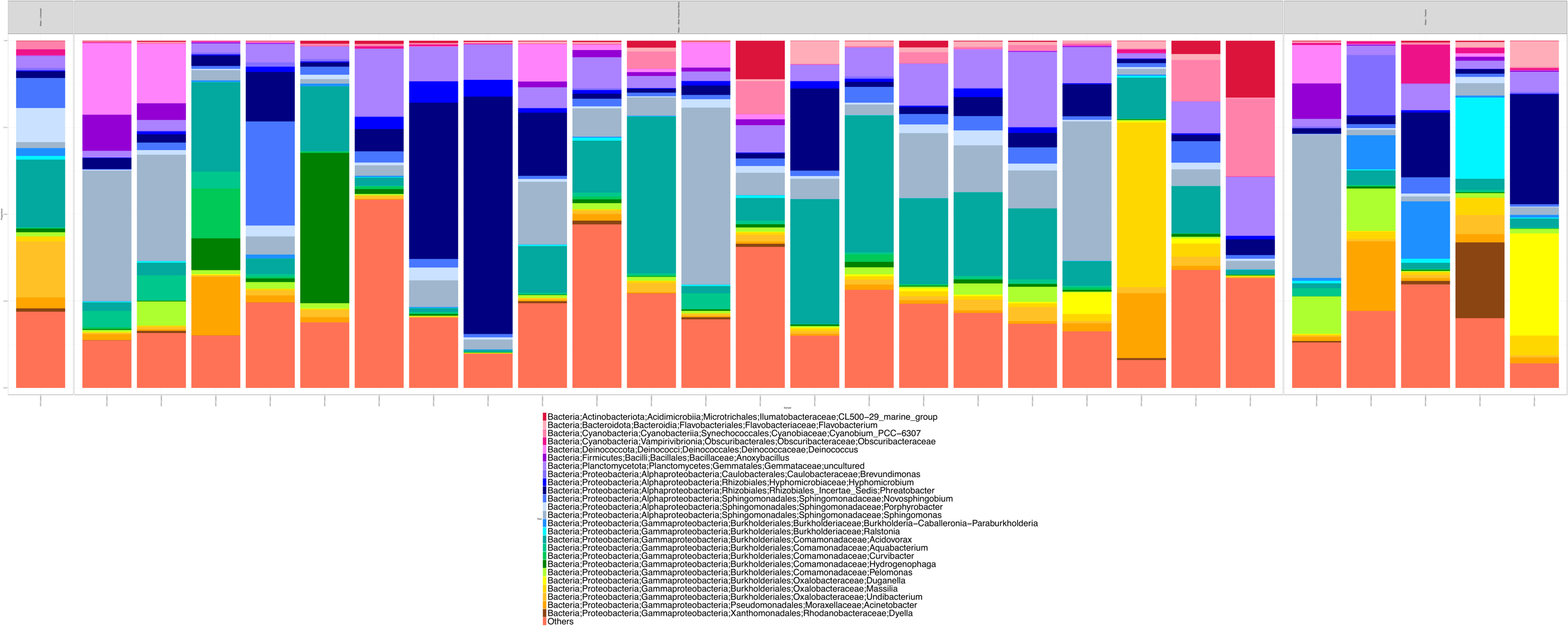

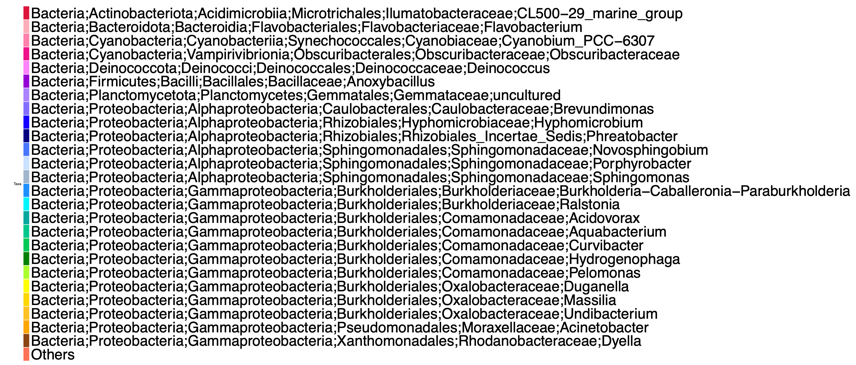

r = 0.9021849 | p = 0.0001 | Sum of Squares = 0.5299 | Correlation = 0.6856 | p = 0.001

### 12. HiSeq V4-V5

a)

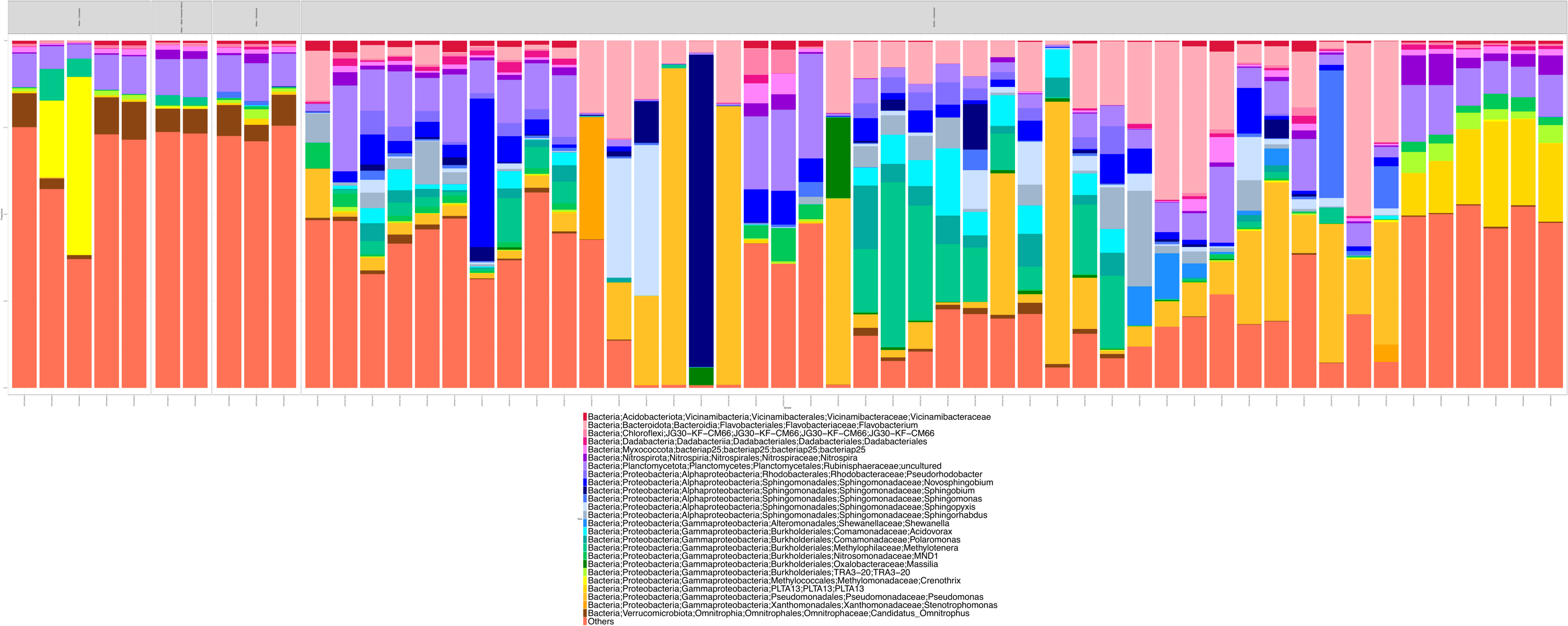

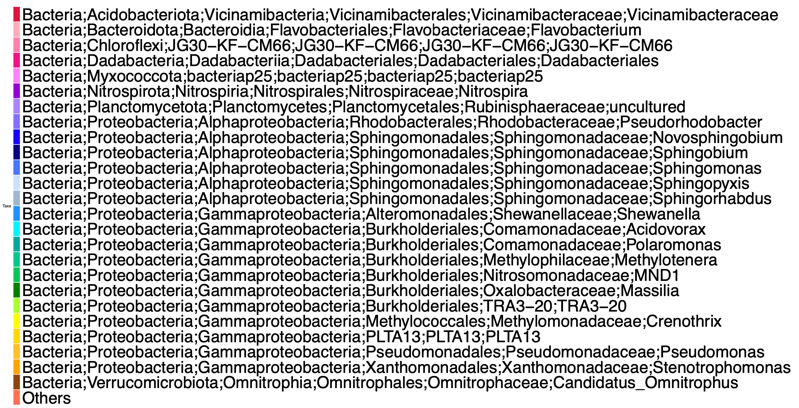

b)

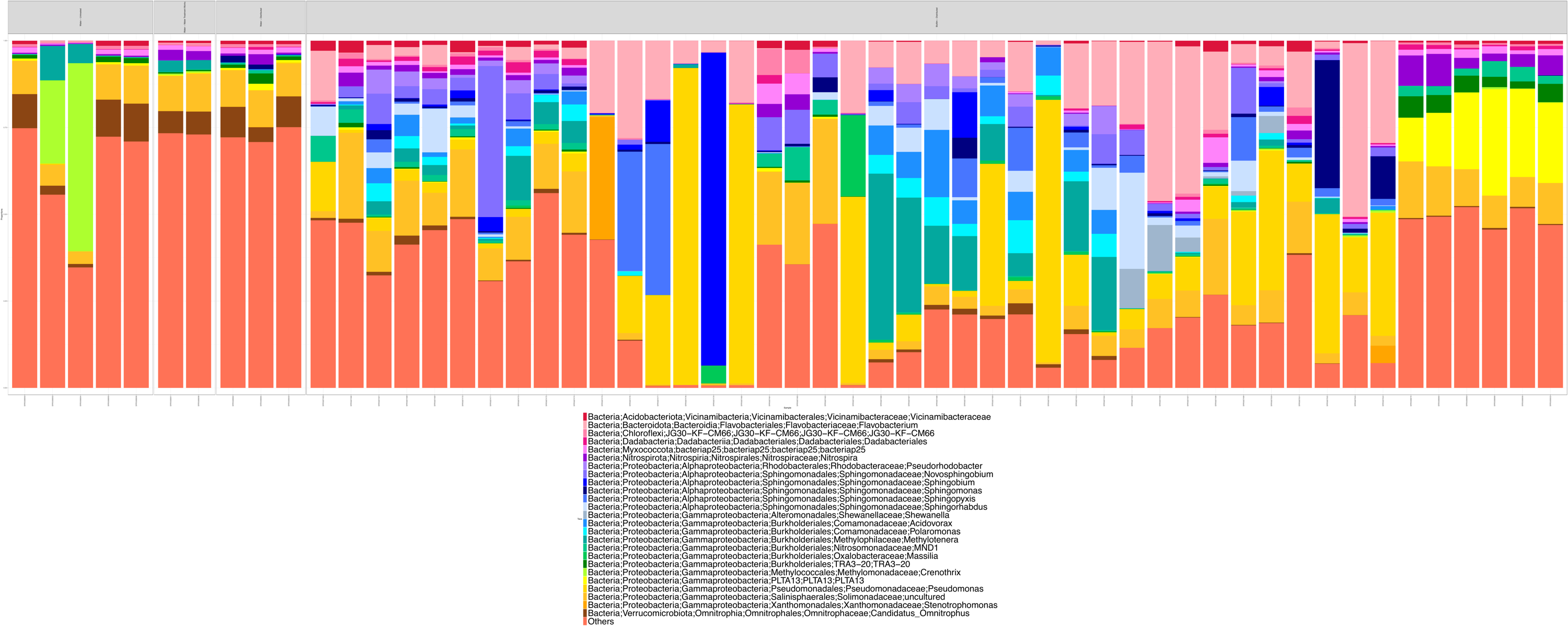

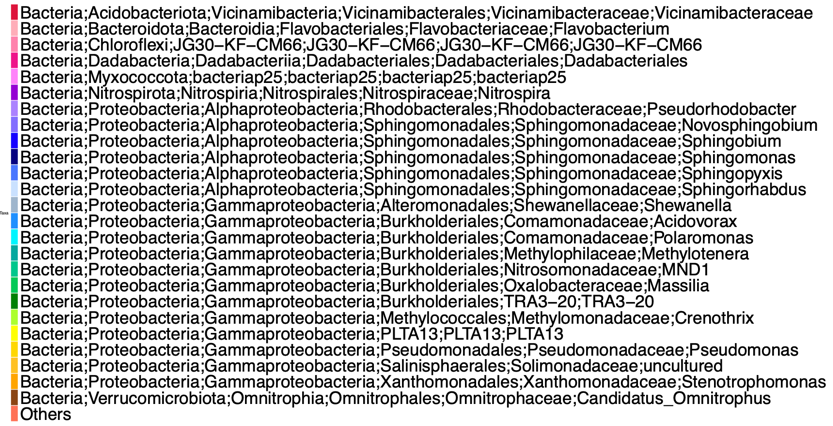

r = 0.7873318| p = 0.0001 | Sum of Squares = 0.4811 | Correlation = 0.7203 | p = 0.001
